# Geminiviral genomes encode additional proteins with specific subcellular localizations and virulence function

**DOI:** 10.1101/2021.03.01.433473

**Authors:** Pan Gong, Huang Tan, Siwen Zhao, Hao Li, Hui Liu, Yu Ma, Xi Zhang, Junjie Rong, Xing Fu, Rosa Lozano-Durán, Fangfang Li, Xueping Zhou

## Abstract

Geminiviruses are plant viruses with limited coding capacity. Geminivirus-encoded proteins were identified applying a 10-kDa arbitrary threshold; however, it is increasingly clear that small proteins play relevant roles in biological systems, which calls for the reconsideration of this criterion. Here, we show that geminiviral genomes contain additional ORFs. Using tomato yellow leaf curl virus, we demonstrate that some of these novel ORFs are expressed during the infection, and that the encoded proteins display specific subcellular localizations. We prove that the largest of these new ORFs, which we name V3, is required for full viral infection, and that the V3 protein localizes in the Golgi apparatus and functions as an RNA silencing suppressor. These results imply that the repertoire of geminiviral proteins can be expanded, and that getting a comprehensive overview of the molecular plant-geminivirus interactions will require the detailed study of small ORFs so far neglected.

## INTRODUCTION

Viruses are intracellular parasites that heavily rely on the host cell machinery to complete their life cycle. Most viruses have small genome sizes, with the concomitant limitation in coding capacity; in order to overcome the restrictions imposed by their reduced proteome, viruses have evolved to encode multifunctional proteins that efficiently target hub proteins in their host cells (reviewed in Brito and Pinney, 2017; King et al., 2018). Nevertheless, higher numbers of virus-encoded proteins might enable more sophisticated infection mechanisms, and therefore maximization of the coding space would be expected to be an advantage to the pathogen.

Geminiviruses are a family of plant viruses with circular, single-stranded (ss) DNA genomes causing devastating diseases in crops around the globe. This family includes nine genera, based on host range, insect vector, and genome structure: *Becurtovirus, Begomovirus, Curtovirus, Eragrovirus, Mastrevirus, Topocuvirus, Turncurtovirus, Capulavirus*, and *Grablovirus* (Zerbini et al., 2017); most species described to date belong to the genus *Begomovirus*. Members of this family have small genomes, composed of one or two DNA molecules of less than 3 Kb each, in which the use of coding space is optimized by bidirectional and partially overlapping open reading frames (ORFs): in one <3 Kb molecule, geminiviruses contain up to 7 ORFs, with a known maximum of 8 viral proteins per species. The geminiviral infection cycle is complex, and multiple steps remain to be fully elucidated. Following transmission by an insect vector, the geminiviral DNA genome must be released from the virion and reach the nucleus, where it will be converted into a double-stranded (ds) DNA replicative intermediate; this dsDNA molecule will serve as template for the transcription of viral genes, including the replication-associated protein (Rep), which reprograms the cell cycle and recruits the host DNA replication machinery. Rolling-circle replication ensues, by which new ssDNA copies of the viral genome are produced. Eventually, the virus must move intracellularly, intercellularly, and systemically, invading new cells and making virions available for acquisition by the vector. In order to accomplish a successful infection, geminiviruses must tailor the cellular environment to favour their replication and spread; for this purpose, they modify the transcriptional landscape of the infected cell, re-direct post-transcriptional modifications, and interfere with hormone signalling, among other things (reviewed in Aguilar et al., 2020; Kumar, 2019; Liu et al., 2021), ultimately suppressing anti-viral defences, creating conditions favourable to viral replication, and manipulating plant development. Although geminivirus-encoded proteins are described as multifunctional, how the plethora of tasks required for a fruitful infection can be performed by only 4-8 proteins is an intriguing biological puzzle. Whether members of this family encode additional small proteins, below the arbitrary 10 kDa threshold established following identification of the first geminivirus species, remains elusive.

Tomato yellow leaf curl virus (TYLCV) is a monopartite begomovirus, causal agent of the destructive tomato leaf curl disease (Basak, 2016). The TYLCV genome contains six known open reading frames (ORFs), encoding the capsid protein (CP)/V1 and V2 in the virion strand, and C1/Rep, C2, C3, and C4 in the complementary strand. Rep creates a cellular environment permissive for viral DNA replication and attracts the DNA replication machinery to the viral genome (reviewed in Hanley-Bowdoin et al., 2013); C2 suppresses post-transcriptional gene silencing (PTGS) (Luna et al., 2012), protein ubiquitination (Lozano-Duran et al., 2011) and jasmonic acid (JA) signalling (Lozano-Duran et al., 2011; Rosas-Diaz et al., 2016); C3 interacts with PCNA, NAC, and the regulatory subunits of DNA polymerases α and δ to enhance viral replication (Castillo et al., 2003; Settlage et al., 2005; Wu et al., 2020); C4 is a symptom determinant, interferes with the intercellular movement of PTGS, and hampers salicylic acid (SA)-dependent defences (Luna et al., 2012; Medina-Puche et al., 2020; Rosas-Diaz et al., 2018); the CP forms the viral capsid and is essential for the transmission by the insect vector, and shuttles the viral DNA between the nucleus and the cytoplasm (Azzam et al., 1994; Diaz-Pendon et al., 2010; Gotz et al., 2012; Kunik et al., 1998; Ohnesorge and Bejarano, 2009; Palanichelvam et al., 1998; Rojas et al., 2001; Rubinstein and Czosnek, 1997); V2 is a strong suppressor of PTGS as well transcriptional gene silencing (TGS), and it mediates the nuclear export of CP (Wang et al., 2014; Wang et al., 2018; Wang et al., 2020; Zhao et al., 2020; Zrachya et al., 2007). Interestingly, a recent report identified 21 transcription initiation sites within the TYLCV genome by taking advantage of cap-snatching by rice stripe virus (RSV) in the experimental *Solanaceae* host *Nicotiana benthamiana*, suggesting that transcripts beyond those encoding these known ORFs might exist (Lin et al., 2017). This idea is further indirectly supported by the fact that attempts at knocking-in tags in geminiviral genomes have so far been fruitless.

Here, we report that geminiviral genomes contain additional ORFs besides the canonical ones described to date. These previously neglected ORFs frequently encode proteins that are phylogenetically conserved. Using the geminivirus TYLCV as an example, we show that some of these ORFs are transcribed during the viral infection, and that the proteins they encode accumulate in the plant cell and show specific subcellular localizations and distinctive features. Moreover, we demonstrate that one of these novel ORFs, which we have named V3 and is conserved in begomoviruses, is essential for full infectivity in *N. benthamiana* and tomato, and encodes a Golgi-localized protein that acts as a suppressor of PTGS and TGS. Taken together, our results indicate that geminiviruses encode additional proteins to the ones described to date, which may largely expand the geminiviral proteome and its intersection with the host cell.

## RESULTS

### The TYLCV genome contains additional conserved open reading frames

It is increasingly clear that small proteins (<100 aa) are prevalent in eukaryotes, including plants, and have biological functions (reviewed in Eguen et al., 2015; Hsu and Benfey, 2018; Murphy et al., 2012). In order to explore whether geminiviral genomes may contain additional ORFs encoding small proteins of predictable functional relevance, we designed a tool that we called ViralORFfinder; this tool uses the ORFfinder from NCBI (https://www.ncbi.nlm.nih.gov/orffinder/) to identify ORFs in an inputted subset of DNA sequences (geminiviral genomes, in this case) and creates a small database with the translated protein sequences, which can be used to BLAST a protein of choice, therefore assessing conservation among the selected species (Figure 1a). The distribution of the protein of interest is then displayed in a phylogenetic tree of the inputted viral species, generated based on the DNA sequences provided. Using ViralORFfinder, additional ORFs can be consistently predicted in geminiviruses of different genera, as illustrated for bipartite begomoviruses (Supplementary figure 1; Supplementary table 2), curtoviruses (Supplementary figure 2; Supplementary table 3), and mastreviruses (Supplementary figure 3; Supplementary table 4); the proteins encoded by some of these ORFs are conserved among species.

**Figure 1.**
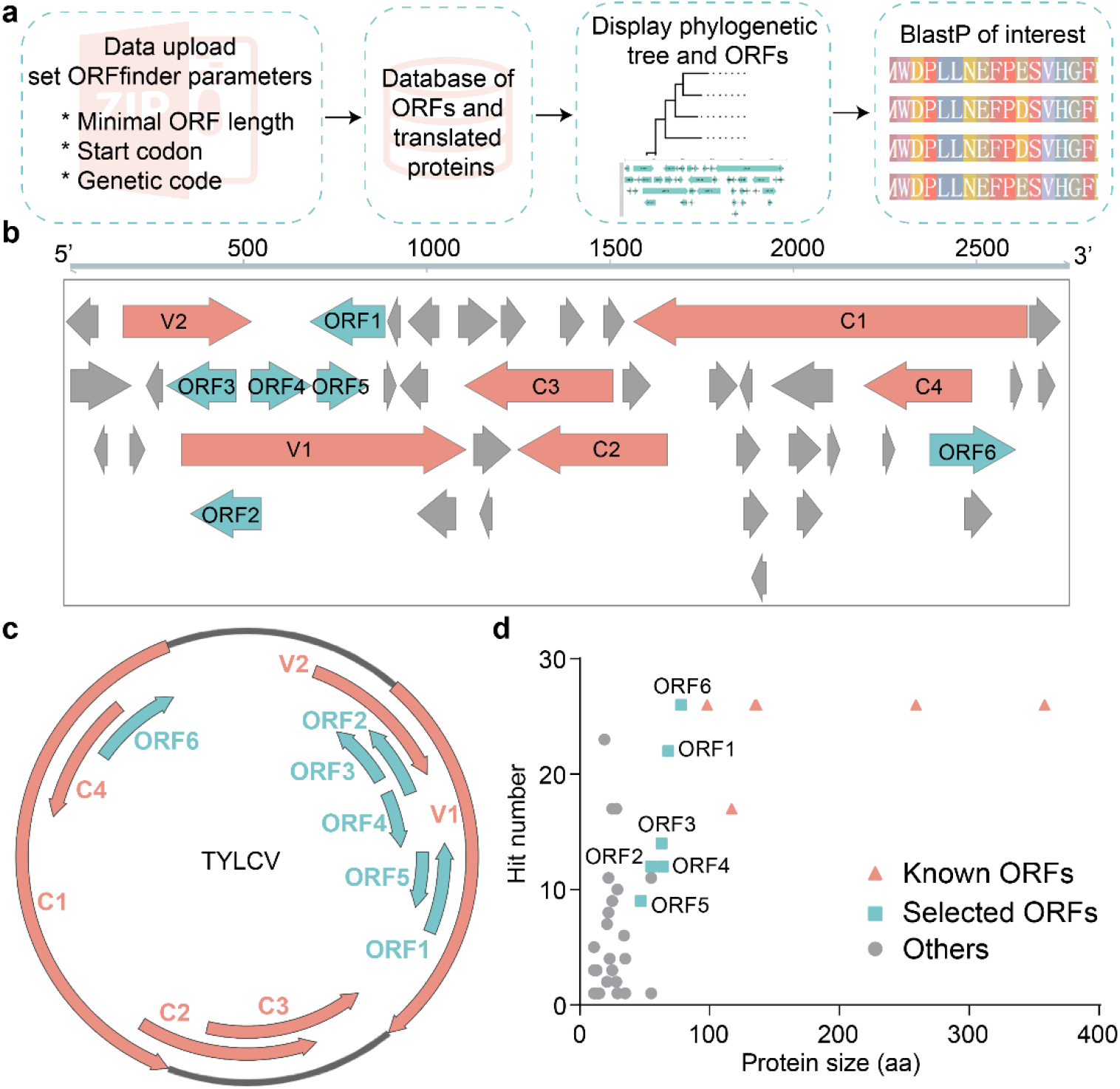
The TYLCV genome contains additional conserved open reading frames. a. Working pipeline of ViralORFfinder, a web-based tool for ORF prediction and protein conservation analysis. b. Schematic view of predicted ORFs (≥ 30 nt) in the TYLCV genome. c. Genome organization of TYLCV; arrows indicate ORFs. In b and c, pink arrows represent the six known ORFs (C1, C2, C3, C4, V1, and V2), while blue arrows represent the six new ORFs described in this work (ORF1-6). d. Correlation between the size of proteins encoded by the ORFs in the TYLCV genome and their representation in the selected subset of begomoviruses (see Supplementary table 6). The TYLCV isolate used in these experiments is TYLCV-Alm.

We then used ViralORFfinder to identify additional proteins encoded by monopartite begomoviruses causing tomato leaf curl disease isolated from different regions of the world (see Methods section; Supplementary figure 4; Supplementary table 5), and identify those present in TYLCV and conserved in other species. As shown in Figure 1b, 43 ORFs encoding proteins of >10 aa were identified in the TYLCV genome. Interestingly, a significant correlation can be found between the size of the encoded proteins and their representation in the selected subset of species, with the six larger proteins (>10 kDa) present in all of them (Supplementary figure 4b, pink). For further analyses, we selected the 6 ORFs that followed in size and prevalence (Supplementary figure 4b, blue), named ORF1-6; the position of these ORFs in the TYLCV genome is shown in Figure 1c. Of note, the proteins encoded by these ORFs are also conserved in other members of the *Begomovirus* genus, both bipartite and monopartite, infecting a broad range of hosts (Supplementary figures 5-7; Supplementary table 6; Figure 1d).

### The novel ORFs in TYLCV encode proteins with predicted domains and specific subcellular localizations

With the aim of gaining insight into the properties of the proteins encoded by the new ORFs from TYLCV, we investigated the presence in their sequence of predicted domains or signals, namely transmembrane domains (TM), nuclear localization signal (NSL), and chloroplast transit peptide (cTP). As shown in Figure 2a, while none of these proteins contains an NLS, and only one of them contains a cTP, three of them contain a predicted TM, which is not present in any of the previously characterized proteins.

**Figure 2.**
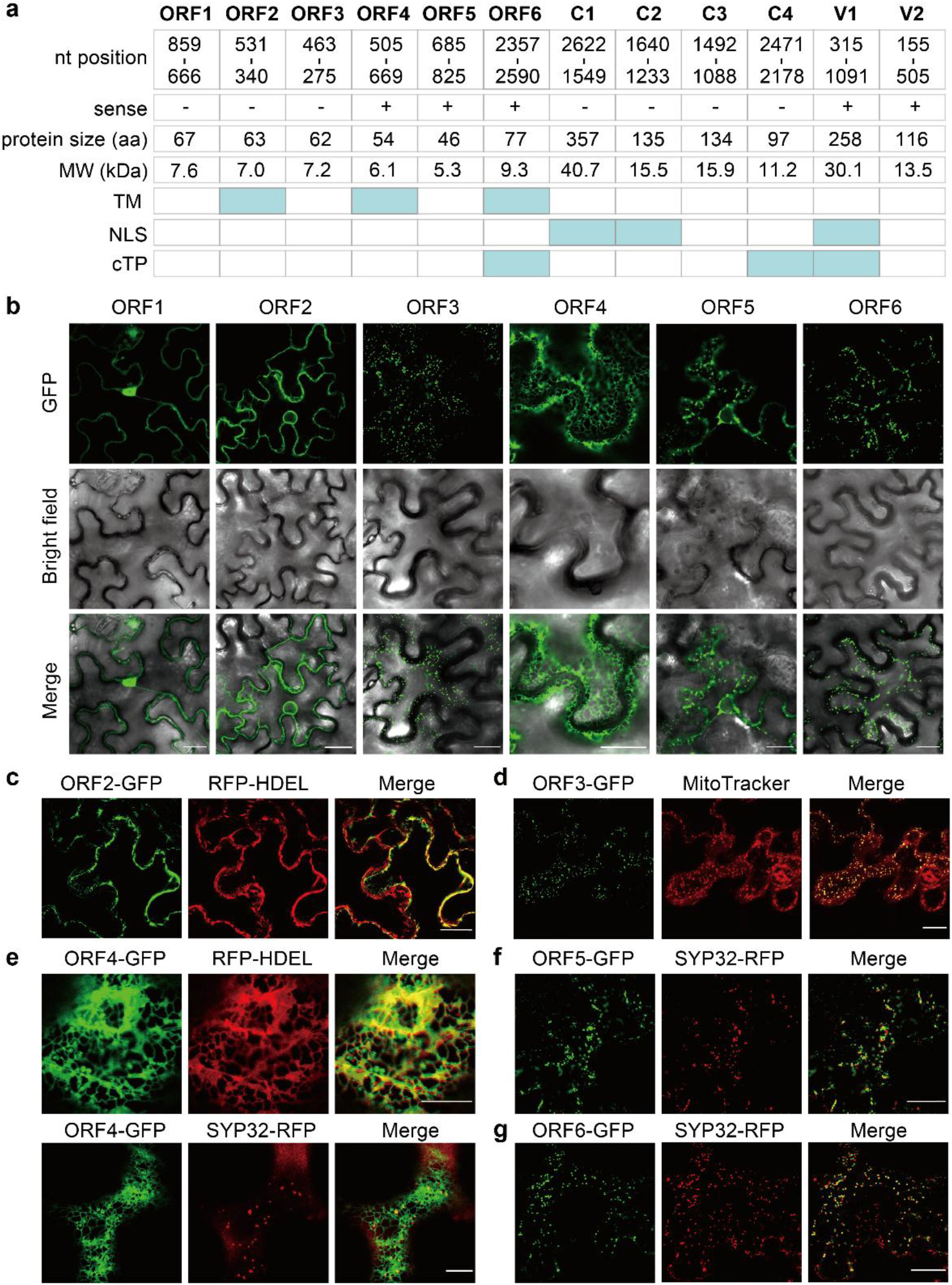
The novel open reading frames in the TYLCV genome encode proteins with predicted domains and specific subcellular localizations. a. Nucleotide position of the six novel ORFs and the six known ORFs in the TYLCV genome, size (in aa) and predicted molecular weight (MW; in kDa) of the corresponding encoded proteins, and domains or signals predicted in the proteins sequence. TM: transmembrane domain; NLS: nuclear localization signal; cTP: chloroplast transit peptide. b. Subcellular localization of the proteins encoded by ORF1-6 fused to GFP at their C-terminus transiently expressed in *N. benthamiana* leaves. Scale bar: 25 μm. c. Co-localization of ORF2-GFP with the ER marker RFP-HDEL. Scale bar: 25 μm. d. Co-localization of ORF3-GFP with the mitochondrial stain MitoTracker Red. Scale bar: 25 μm. e. Co-localization of ORF4-GFP with the ER marker RFP-HDEL and the cis-Golgi marker SYP32-RFP. Scale bar: 10 μm. f. Co-localization of ORF5-GFP with the cis-Golgi marker SYP32-RFP. Scale bar: 25 μm. g. Co-localization of ORF6-GFP with the cis-Golgi marker SYP32-RFP. Scale bar: 25 μm. b-g. These experiments were repeated at least three times with similar results; representative images are shown. The TYLCV isolate used in these experiments is TYLCV-Alm.

We then cloned ORF1-6, fused them to the GFP gene, and transiently expressed them in *N. benthamiana* leaves; confocal microscopy indicates that these fusion proteins present specific subcellular localizations (Figure 2b). Whereas the ORF1-encoded protein is mostly nuclear, co-expression with marker proteins or dyes unveils that the ORF2-encoded protein localizes in the endoplasmic reticulum (ER), as demonstrated by the co-localization with RFP-HDEL; the ORF3-encoded protein in mitochondria, as demonstrated by the co-localization with MitoTracker; the ORF4-encoded protein in the ER and the Golgi apparatus, as demonstrated by the partial co-localization with RFP-HDEL and SYP32-RFP; and the ORF5- and ORF6-encoded proteins mostly in Golgi, as demonstrated by the partial co-localization with SYP32-RFP (Figure 2c-g; Supplementary figure 8). The specific subcellular localization exhibited by each of these proteins suggests that, despite their small size (5.3-9.3 kDa; Figure 2a), either their sequence contains the appropriate targeting signals, or they interact with plant proteins that enable their precise targeting in the cell.

A prerequisite for these ORFs to have a biological function is their expression in the context of the viral infection. Therefore, we cloned the TYLCV genomic sequences upstream of each ATG (pORF1-6) before the GFP reporter gene and tested their promoter activity in transiently transformed *N. benthamiana* leaves. As shown in Supplementary figure 9, none of these sequences could drive GFP expression in the absence of the virus, but pORF1, pORF2, pORF4, and pORF5 could when the virus was present; the sequence upstream of the C4 ORF was used as positive control, and could activate GFP expression both in the presence and absence of TYLCV. The promoter activity of these upstream sequences in infected cells strongly suggests that at least ORF1, 2, 4, and 5 are expressed during the viral infection.

### The novel V3 protein from TYLCV is a Golgi-localized silencing suppressor required for full infection

ORF6 is the largest of the newly described ORFs in the TYLCV genome, and the protein it encodes displays the highest degree of conservation in a selected subset of 26 representative begomoviruses (Supplementary figure 7). Therefore, we decided to further characterize this ORF as a proof-of-concept of the potential biological roles of novel ORFs. Hereafter, ORF6 will be referred to as V3, since it is the third ORF on the viral strand in the TYLCV genome.

The V3 ORF is located in positions 2350-2583 of the TYLCV genome (TYLCV-BJ; Figure 1d), and encodes a 77-amino acid protein. In different begomovirus species, the V3 ORF ranges from 87 to 234 nt, and the protein it encodes presents a high degree of similarity, with 6 residues completely conserved (Supplementary figure 7).

In order to determine whether V3 is transcribed during the viral infection, we checked if the corresponding transcript was present in TYLCV-infected samples: as shown in Figure 3a, the V3 transcript was found upon TYLCV infection, but not in uninfected plants. 5’ rapid amplification of cDNA ends (RACE) was then used to identify the transcriptional initiation site of V3, which was found to be located between 176 and 421 upstream of the start codon (Figure 3b). Interestingly, most of the sites identified by RACE (7 out of 11) are close to A2058, which was previously isolated by cap-snatching of RSV (Lin et al., 2017).

**Figure 3.**
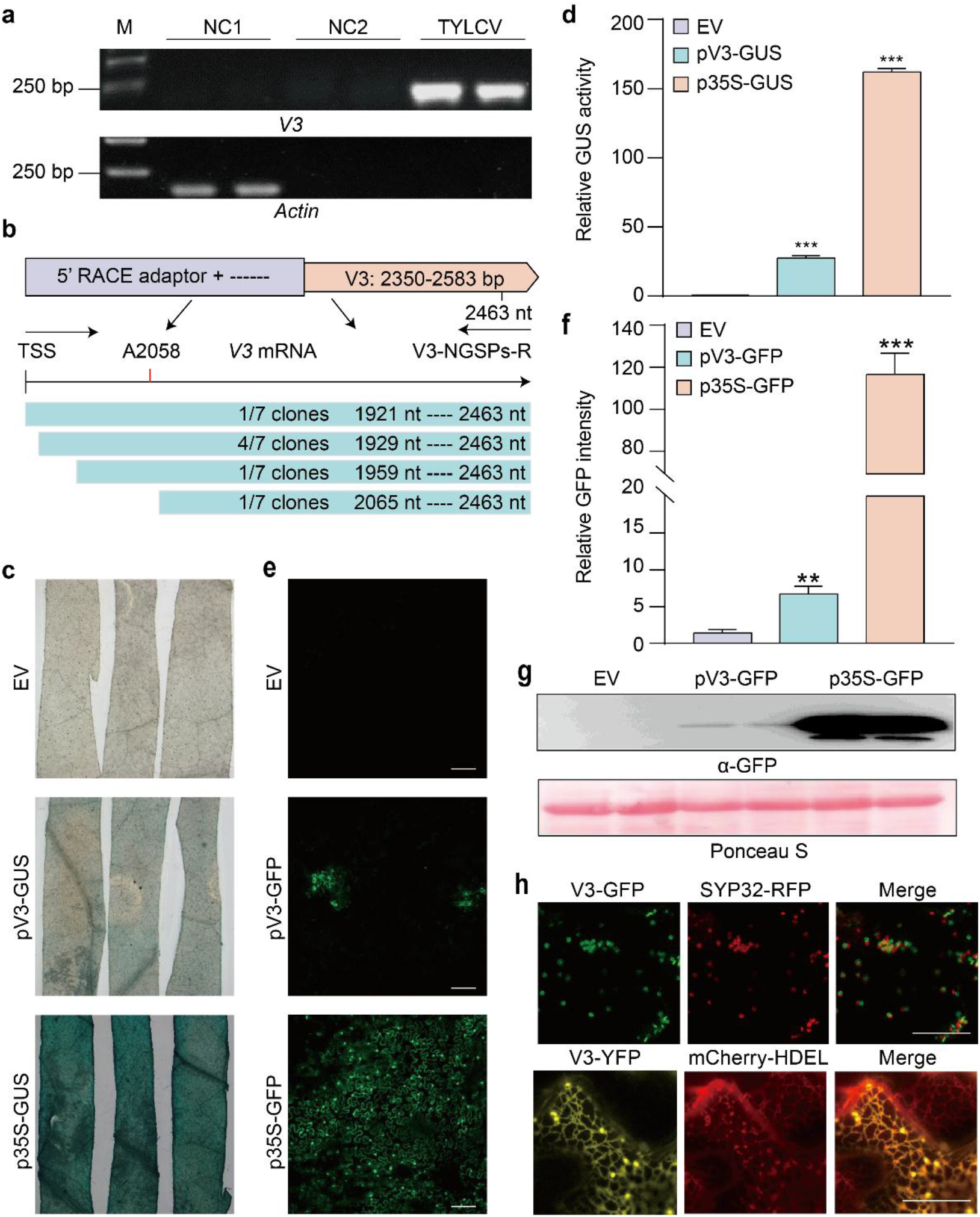
ORF6/V3 is expressed during the infection and encodes a Golgi-localized protein. a. RT-PCR analysis of V3 transcripts from TYLCV-infected or uninfected *N. benthamiana* plants. M: DNA ladder marker. NC1: negative control 1 (reverse-transcription of total RNA extracted from uninfected plants with RT Primers). NC2: negative control 2 (reverse-transcription of total RNA extracted from uninfected plants with V3-specific primers). TYLCV: reverse-transcription of total RNA extracted from TYLCV-infected plants with V3-specific primers. b. Transcriptional start site analysis of TYLCV V3 by 5’ RACE. TSS: transcription start site. A2058: V3 TSS captured by RSV cap-snatching (Lin et al., 2017). c. Activity of pV3 promoter (and p35S promoter as positive control) in promoter-GUS fusions in transiently transformed *N. benthamiana* leaves at 2 dpi. EV: empty vector. d. Quantification of relative GUS activity in samples from (c). Error bars represent SD of n=3. Asterisks indicate a statistically significant difference according to Student’s t-test, *** p < 0.001. e. Activity of pV3 promoter (and p35S promoter as positive control) in promoter-GFP fusions in transiently transformed *N. benthamiana* leaves at 2 dpi. Scale bar: 100 μm. EV: empty vector. f. Quantification of relative GFP intensity in samples from (e). Error bars represent SD of n=3. Asterisks indicate a statistically significant difference according to Student’s t-test, ** p < 0.01, *** p < 0.001. g. Western blot analysis of GFP protein from (e). Ponceau S staining of the large RuBisCO subunit serves as loading control. EV: empty vector. h. Co-localization of V3-GFP with the cis-Golgi maker SYP32-RFP (upper panel) and co-localization of V3-YFP with the ER maker mCherry-HDEL (lower panel). Scale bar: 20 μm. The TYLCV isolate used in these experiments is TYLCV-BJ.

The finding that a transcript corresponding to the V3 ORF can be identified in infected samples strongly suggests that the sequence upstream of this ORF must act as a promoter. Since the 500 bp fragment previously tested (Supplementary figure 9) did not show promoter activity, we tested a larger, 833-nt sequence upstream of the V3 start codon, which was cloned before the GUS or GFP reporter genes. As shown in Figure 3c, d, this viral sequence could activate GUS expression, leading to detectable GUS activity, and it could also drive expression of GFP (Figure 3e-g), albeit more weakly than the 35S promoter. Taken together, these results confirm that the V3 ORF can be expressed *in planta* from its genomic context.

Given that protein function is tightly linked to its spatial location, we then decided to analyse the subcellular localization of the V3 protein in detail. As can be seen in Figure 3h, V3 is a Golgi-localized protein; this localization does not change in the presence of the virus (Supplementary figure 10a). A closer observation confirms that V3 is localized to cis-Golgi, as shown by the co-localization of V3-GFP with the markers Man49-mCherry and SYP32-RFP (Figure 3h; Supplementary figure 10b); nevertheless, the YFP fusions YFP-V3 and V3-YFP can also partially co-localize with the endoplasmic reticulum (ER), as indicated by their partial co-localization with the ER marker mCherry-HDEL or RFP-HDEL (Figure 3h; Supplementary figure 10c, d), which may reflect the transition of the protein from the ER to the Golgi apparatus.

With the aim of assessing the biological relevance of the V3 protein for the TYLCV infection, we next generated a mutated infectious clone carrying a T2351C substitution in the V3 ATG, hence impairing the production of the V3 protein. Since the V3 ORF overlaps with the Rep/C1 and C4 ORFs, nt replacements in the start codon of the V3 ORF necessarily affect the protein sequence of the resulting Rep or C4 proteins; the chosen change results in a I89V substitution in the Rep/C1 protein, with no change in C4. This mutant infectious clone is hereafter referred to as TYLCV-mV3.

TYLCV-mV3 was then inoculated into *N. benthamiana* plants, and its performance compared to that of the wild-type (WT) virus (TYLCV-WT). At 10 days post-inoculation (dpi), TYLCV-mV3-infected plants displayed mild leaf curling symptoms and presented lower viral DNA load compared to plants inoculated with TYLCV-WT (Figure 4a, b), which correlated with a lower accumulation of CP (Figure 4c). These differences apparently result from a delay in the infection, measured as symptom appearance (Figure 4d). To evaluate the potential functional impact of the I89V substitution in Rep/C1 on the viral infection and disentangle this effect to that derived from the absence of V3, transgenic *N. benthamiana* lines expressing V3-YFP under a 35S promoter were generated, and a complementation assay was performed. Of note, the V3-expressing plants do not display obvious developmental abnormalities (Supplementary figure 11a); the expression of V3-YFP was confirmed by qRT-PCR and western blot (Supplementary figure 11b, c). As shown in Figure 4e and Supplementary figure 11d, transgenic expression of V3-YFP could fully complement the lack of V3 in the TYLCV-mV3 clone, measured as incidence and severity of symptom appearance, indicating that the lower progression of the infection observed upon inoculation with the V3 null mutant is due to the lack of this protein, and not to a suboptimal performance of Rep-I89V. Next, we evaluated the virulence of TYLCV-mV3 on tomato, the virus’ natural host. As previously observed in *N. benthamiana*, the lack of V3 resulted in lower viral load and CP accumulation, and milder symptoms at 10 dpi (Figure 4f-h), confirming that V3 plays a relevant role in the viral infection that is not restricted to *N. benthamiana*.

**Figure 4.**
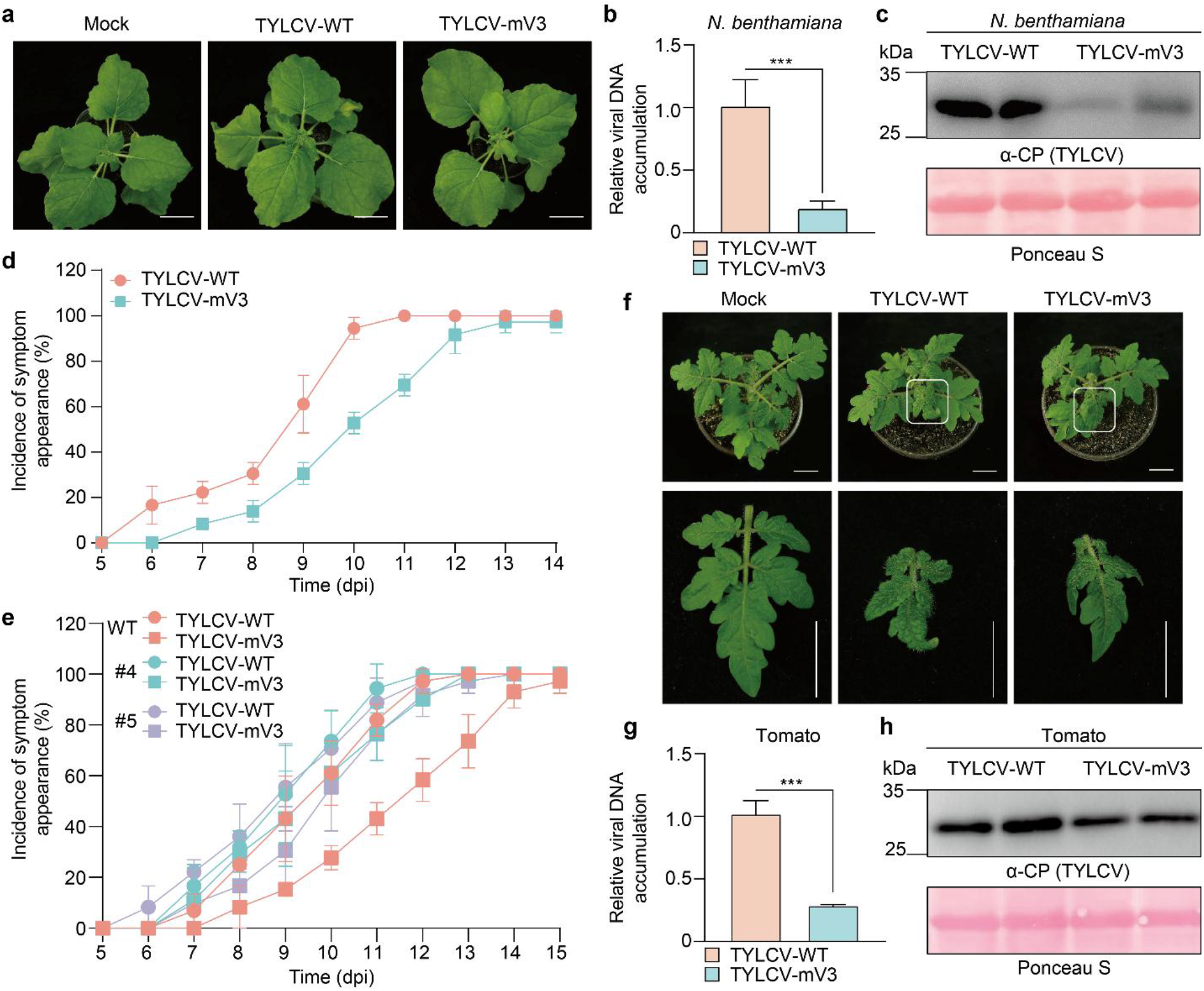
V3 is required for full TYLCV infection in *N. bentamiana* and tomato. a. Symptoms of *N. benthamiana* plants inoculated with wild-type TYLCV (TYLCV-WT), a V3 null mutant TYLCV (TYLCV-mV3), or mock-inoculated (pCAMBIA2300 empty vector), at 10 dpi. Bar = 2 cm. b. Viral DNA accumulation in TYLCV-WT- and TYLCV-mV3-infected plants in (a), measured by qPCR. Error bars represent means ± SD of n=3. Asterisks indicate a statistically significant difference according to Student’s t-test, *** p<0.001. 25S RNA was used as internal reference. c. Western blot showing TYLCV CP accumulation in systemic leaves of TYLCV-WT- and TYLCV-mV3-infected plants from (a). Ponceau S staining of the large RuBisCO subunit serves as loading control. d, e. Incidence of symptom appearance in WT (d, e) or V3 transgenic (e) *N. benthamiana* plants infected with TYLCV-WT or TYLCV-mV3. Error bars represent means ± SD from three independent experiments; at least 8 plants were used per viral genotype and experiment. For images of symptoms in the V3 transgenic lines, see Supplementary figure 10d. f. Symptoms of tomato plants inoculated with WT TYLCV (TYLCV-WT), a V3 null mutant TYLCV (TYLCV-mV3), or mock-inoculated (pCAMBIA2300 empty vector), at 10 dpi. Bar = 2 cm. g. Viral DNA accumulation in TYLCV-WT- and TYLCV-mV3-infected plants from (f), measured by qPCR. Error bars represent means ± SD of n=3. Asterisks indicate a statistically significant difference according to Student’s t-test, *** p<0.001. 25S RNA was used as internal reference. h. Western blot showing TYLCV CP accumulation in systemic leaves of TYLCV-WT- and TYLCV-mV3-infected plants from (f). Ponceau S staining of the large RuBisCO subunit serves as loading control. The TYLCV isolate used in these experiments is TYLCV-BJ.

Heterologous expression from a potato virus X (PVX)-derived vector and quantification of the impact on PVX pathogenicity is a widely used approach to test virulence activity of viral genes of interest. Confirming a contribution of V3 to virulence, the presence of this gene in PVX-V3 led to an exacerbation of disease symptoms compared to PVX alone at 10 and 30 dpi, with a concomitant higher accumulation of the PVX CP at 30 dpi (Supplementary figure 12); βC1, a symptom determinant encoded by tomato yellow leaf curl China betasatelite, was used as positive control. Infection by PVX-V3, however, did not lead to H2O2 accumulation or cell death (Supplementary figure 12c).

It has been previously established that a high correlation exists between the ability of a viral protein to suppress RNA silencing and its capacity to enhance the severity of the PVX infection. RNA silencing is conserved in eukaryotes, and is considered the main anti-viral defence mechanism in plants (Ding, 2010; Ding et al., 2004; Jin et al., 2020). Supporting this notion, virtually all plant viruses described to date encode at least one protein with RNA silencing suppression activity (Burgyan and Havelda, 2011; Jin et al., 2020). With the aim to test if V3 can suppress post-transcriptional gene silencing (PTGS), we transiently expressed GFP from a 35S promoter in leaves of transgenic 16c *N. benthamiana* plants, harbouring a 35S:GFP cassette (Ruiz et al., 1998), in the presence or absence of V3; the well-described silencing suppressor P19 from tomato bushy stunt virus was used as positive control. At 4 dpi, fluorescence had already substantially decreased when no viral protein was co-expressed, but was maintained in the samples with P19 or Myc-V3 (Figure 5a). Western blot and qRT-PCR were used to confirm that both the GFP protein as well as the corresponding mRNA accumulated to higher levels in tissues expressing P19 or Myc-V3 (Figure 5b). At 20 dpi, systemic leaves of 16c plants inoculated with the constructs to express either of the viral proteins remained green, while fluorescence had disappeared in control plants as a result of systemic silencing (Figure 5a).

**Figure 5.**
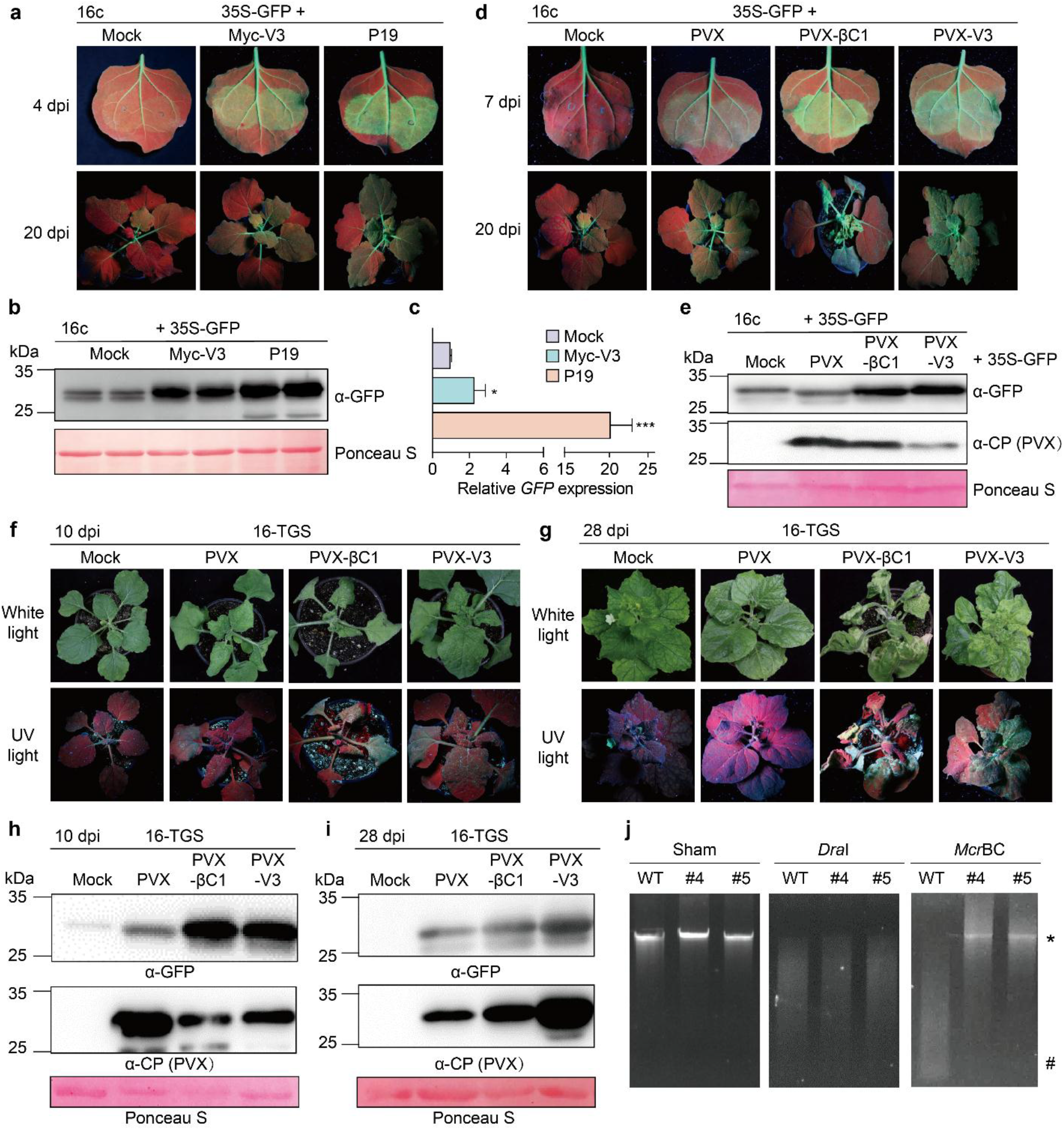
V3 functions as a suppressor of PTGS and TGS. a. Transgenic 16c *N. benthamiana* plants co-infiltrated with constructs to express GFP (35S-GFP) and Myc-V3, P19 (as positive control), or mock (empty vector, as negative control) at 4 dpi (upper panel) or 20 dpi (lower panel) under UV light. b. Western blot showing the GFP accumulation in inoculated leaves from (a) at 4 dpi. The corresponding Ponceau S staining of the large RuBisCO subunit serves as a loading control. c. Relative GFP mRNA accumulation in inoculated leaves from (a) at 4 dpi measured by qRT-PCR. Error bars represent means ± SD of n=3. Asterisks indicate a statistically significant difference according to Student’s t-test, * p<0.5, *** p<0.001. *NbActin2* was used as the internal reference. d. Transgenic 16c *N. benthamiana* plants co-infiltrated with constructs to express GFP (35S-GFP) and PVX, PVX-V3, PVX-βC1 (as positive control), or mock (infiltration buffer, as negative control) at 7 dpi (upper panel) or 20 dpi (lower panel) under UV light. e. Western blot showing the GFP accumulation in inoculated leaves from (d) at 7 dpi. The corresponding Ponceau S staining of the large RuBisCO subunit serves as a loading control. f, g. Symptoms of 16-TGS *N*. *benthamiana* plants infected with PVX, PVX-V3, PVX-βC1 (as positive control), or mock-inoculated under white light or UV light at 10 dpi (f) and 28 dpi (g). h. i. Western blot showing accumulation of GFP and PVX CP in systemically infected leaves from (f) and (i). The corresponding Ponceau S staining of the large RuBisCO subunit serves as loading control. j. DNA methylation analysis by restriction enzyme digestion in V3-YFP transgenic *N. benthamiana* plants. Genomic DNA extracted from WT *N. benthamiana* or two independent V3-YFP transgenic lines (#4 and #5) was digested with the methylation-dependent restriction enzyme *McrBC* and the methylation-insensitive enzyme *Dra*I. ‘Sham’ indicates a mock digestion with no enzyme added. The positions of undigested input genomic DNA is indicated with an asterisk; the position of the *Mcr*BC-digested products is indicated with a hashtag.

The ability of V3 to suppress PTGS was further confirmed by expressing this protein from a PVX-based vector; in this case, βC1, which also functions as silencing suppressor, was used as positive control. Transient co-transformation of a PVX infectious clone together with a 35S:GFP cassette in leaves of 16c plants led to weak fluorescence in the infiltrated tissues at 7 dpi, and systemic silencing at 20 dpi; in stark contrast, co-transformation with PVX-βC1 or PVX-V3 resulted in the maintenance of strong fluorescent signal at 7 dpi, and absence of systemic silencing (Figure 5d). Neither βC1 nor V3 enhanced the local accumulation of PVX, as indicated by the accumulation of the PVX CP protein (Figure 5e), hence ruling out an indirect effect of these proteins on the endogenous ability of PVX to suppress silencing. Therefore, our results demonstrate that V3 from TYLCV can effectively suppress PTGS in plants.

Another level of RNA silencing is transcriptional gene silencing (TGS), which acts through methylation of DNA at cytosine residues; TGS also acts as an anti-viral response in plants, and is particularly relevant against geminiviruses, which replicate their DNA genomes in the nucleus of the infected cell (Jin et al., 2020; Wang et al., 2019). To investigate whether V3 can also act as a TGS suppressor, we inoculated 16-TGS plants, in which the GFP transgene is silenced due to methylation of the 35S promoter (Yang et al., 2011), with PVX or PVX-V3; PVX-βC1 was used as positive control, since βC1 can also act as a TGS suppressor. At 10 dpi, green fluorescence could be observed in systemic leaves of 16-TGS plants inoculated with PVX-βC1 and PVX-V3, as opposed to mock- or PVX-inoculated plants (Figure 5f); this fluorescence persisted at 28 dpi (Figure 5g). Visual assessment was confirmed at the molecular level by western blot (Figure 5h, i), indicating that V3 can also suppress TGS in the host plant.

To confirm the effect of V3 on DNA methylation, the level of genome-wide methylation in transgenic V3-YFP plants was examined by digestion with a methylation-dependent restriction enzyme, *McrBC* (Stewart et al., 2000). As presented in Figure 5j, genomic DNA from two independent V3-YFP lines was completely digested by the methylation-independent restriction endonuclease *Dra*I, but only partially digested by *Mcr*BC, in sharp contrast to the genomic DNA from WT plants. This indicates a lower level of DNA methylation in transgenic plants expressing V3, further supporting a function of this viral protein as TGS suppressor.

In summary, our results demonstrate that V3 is a newly described protein encoded by TYLCV, which preponderantly localizes in the cis-Golgi, significantly contributes to virulence, and functions as a suppressor of both PTGS and TGS.

## DISCUSSION

Geminivirus-encoded proteins have so far been identified taking into consideration an arbitrary threshold of 10 kDa, below which potential proteins were discarded. However, it is increasingly clear that small proteins and peptides play relevant roles in biological systems, hence calling for the reconsideration of this criterion. Here, we show that geminiviral genomes contain additional ORFs beyond those previously described, at least some of which are conserved within members of a given genus, hinting at functional relevance. Using TYLCV as a model, we demonstrate that at least some of these conserved novel ORFs are expressed during the infection, and that the proteins they encode localize in specific subcellular compartments. Interestingly, in this subset of selected proteins novel localizations, not described for any of the other TYLCV-encoded proteins, are represented, including the Golgi apparatus (ORF4, 5, and 6) and mitochondria (ORF 3). Also of note, some of these proteins (ORF2, 4, and 6) harbor transmembrane domains, which are not present in any of the “canonical” proteins from TYLCV. These results indicate that the repertoire of geminiviral proteins can be expanded, and that the new additions to the viral proteomes will most likely perform additional virulence functions and/or employ alternative molecular mechanisms to those exhibited by the previously described proteins. We might still be, therefore, far from getting a comprehensive overview of the plant-geminivirus molecular interaction landscape, which will require the detailed study of potentially multiple small ORFs that have so far been neglected.

In this work, we selected the largest of the new ORFs found in TYLCV, which we name V3, to be used as a proof-of-concept of the potential functionality of novel viral small proteins. Strikingly, the lack of V3 negatively impacted the viral infection in two different hosts, *N. benthamiana* and tomato, demonstrating that V3 has a biological function. However, V3 is not essential, since a V3 null mutant virus can still accumulate and establish a systemic infection.

The V3 protein is mostly localized in the cis-Golgi, a novel localization for a geminivirus-encoded protein. Remarkably, V3 can suppress both PTGS and TGS, an ability that may underlie its virulence-promoting effect on TYLCV and PVX. One intriguing question is how V3 can exert this effect from the Golgi apparatus. Interestingly, connections have been drawn between the endomembrane system and RNA silencing (reviewed in Kim et al., 2014). In the model plant *Arabidopsis thaliana*, electron microscopy unveiled an enrichment of the Argonaute protein AGO1, a central player in PTGS, in close proximity to Golgi (Derrien et al., 2012); another Argonaute protein, AGO7, which has been recently shown to play a role in anti-viral defence (Zheng et al., 2019), also co-purifies with membranes and concentrates in cytoplasmic bodies linked to the ER/Golgi endomembrane system (Jouannet et al., 2012). It seems therefore plausible that this subcellular localization is permissive for a direct targeting of RNA silencing, although further experiments will be necessary to uncover the exact molecular mechanism underlying this activity of V3.

PTGS and TGS are arguably the main plant defence mechanisms against geminiviruses. This idea is supported by the fact that, despite limited coding capacity, a given geminivirus species can produce several proteins that independently target these processes. TYLCV encodes at least three proteins capable of acting as PTGS suppressors, namely C2, C4, and V2 (Luna et al., 2012; Rosas-Diaz et al., 2018; Zrachya et al., 2007), and at least two, Rep and V2, capable of suppressing TGS (Rodriguez-Negrete et al., 2013; Wang et al., 2014; Wang et al., 2018; Wang et al., 2020); these proteins exert their functions through non-overlapping mechanisms. Only one of the viral proteins, V2, has been described as a simultaneous suppressor of PTGS and TGS, as observed for V3. All of these silencing suppressors encoded by TYLCV are essential for the infection, although this may be due to their contribution to additional virulence activities, enabled by their multifunctional nature. Similarly, it is possible that V3 exerts additional, yet-to-be described functions during the viral infection.

Why a given geminivirus species needs multiple proteins targeting the same pathway is a thought-provoking question. Since the viral infection is a process, the temporal dimension must be considered: geminiviral genes can be classified as early or late, depending on the timing of their expression, with the strongest PTGS and TGS suppressor, V2, being a late gene, probably due to the requirement of another viral protein, C2, to activate its expression. The V3 promoter was active in the absence of the infection, which suggests that it can be expressed as an early gene. The early expression of V3 would guarantee the availability of a PTGS and TGS suppressor during the time between the synthesis of the dsDNA replicative intermediate and the expression of V2 later in the cycle, enhancing the effectiveness of viral accumulation and spread. Nevertheless, further work will be required to acquire a full understanding of the potential breadth of functions exerted by V3 and of the underpinning molecular mechanisms.

## METHODS

### Plant materials

*N. benthamiana* and *Solanum lycopersicum* (tomato) plants were grown in a growth chamber with 60% relative humidity and a 16 h:8 h light:dark, 25°C:18°C regime. The transgenic GFP 16c line was kindly provided by David C. Baulcombe (University of Cambridge, UK) (Ruiz et al., 1998); 16-TGS plants were described previously (Buchmann et al., 2009); the transgenic RFP-H2B line was kindly shared by Michael M. Goodin (University of Kentucky, USA) (Martin et al., 2009).

The *Agrobacterium tumefaciens* strain EHA105 containing the pEarleygate101:V3-YFP construct was used to generate 35S: *V3-YFP* transgenic *N. benthamiana* lines by leaf disc transformation as previously described (Li et al., 2015).

### Agroinfiltration and viral inoculation

For *A. tumefaciens-mediated* transient expression in *N. benthamiana*, plasmids were transformed into the EHA105 strain by the freeze-thaw method (Figures 3a-h lower panel, 4, 5, Supplementary figure 10b, c, Supplementary figure 12) or to the GV3101 strain through electroporation (Figures 2, 3h upper panel, Supplementary figure 8, Supplementary figure 9, Supplementary figure 10a, d). *Agrobacterium* cultures were resuspended in infiltration buffer (10 mM MgCl2, 10 mM MES (pH 5.6), and 100 μM acetosyringone) to an OD600 = 0.1-0.5, then infiltrated into the adaxial side of four-week-old *N. benthamiana* leaves with a needle-less syringe. For viral inoculation, two-week-old *N. benthamiana* plants or tomato plants at the two-leaf stage were infiltrated with *Agrobacterium* cultures carrying the TYLCV-BJ (MN432609) infectious clone. For experiments that required co-infiltration, *Agrobacterium* suspensions carrying different constructs were mixed at 1:1 ratio before infiltration.

### Plasmid construction

Viral open reading frames (ORFs) from the TYLCV (TYLCV-Alm, Accession No. AJ489258) genome were cloned in the pENTR^TM^/D-TOPO^®^ vector (Thermo Scientific) (for ORF1, ORF2, ORF4, ORF5, ORF6/V3) or the pDONR^TM^/Zeo vector (Thermo Scientific) (for ORF3) without a stop codon. The binary plasmids to express GFP-fused viral proteins were generated by sub-cloning (Gateway LR reaction, Thermo Scientific) the viral ORFs from the corresponding entry vectors into pGWB505 (Nakagawa et al., 2007). To generate the constructs to express V3-YFP, YFP-V3, or Myc-V3, the full-length V3 ORF was obtained from TYLCV (TYLCV-BJ, Accession No. MN432609) and recombined into the binary destination vectors pEarleygate101, pEarleygate104, or pEarleygate203, respectively (Earley et al., 2006). Please note that the V3/ORF6 protein sequences from TYLCV-Alm and TYLCV-BJ are identical.

To generate the construct to express SYP32-RFP (as a cis-Golgi marker), the gene encoding SYNTAXIN OF PLANTS 32 (SYP32) was amplified from *A. thaliana* cDNA, cloned into the pENTR^TM^/D-TOPO^®^ vector (Thermo Scientific), and sub-cloned into pGWB554 (Nakagawa et al., 2007) using a Gateway LR reaction (Thermo Scientific). The construct to express RFP-HDEL is from Liu et al., 2015, and the ones to express mCherry-HDEL and Man49-mCherry are from Nelson et al., 2007.

To generate the constructs used for the analysis of promoter activity, the 500-nt sequence upstream of the ORF1, ORF2, ORF4, ORF5, ORF6/V3, and C4 ATG, or the 570-nt sequence upstream of the ORF3 ATG, were PCR-amplified and cloned into the pDONR^TM^/Zeo vector (Thermo Scientific). These sequences were then sub-cloned into pGWB504 (Nakagawa et al., 2007) by a Gateway LR reaction to generate pORF-GFP. In addition, the 833-nt sequence upstream of the V3/ORF6 ATG was PCR-amplified and cloned into pINT121-GUS digested with *HindIII* and *Bam*HI to generate pINT121-V3-GUS (pV3-GUS), or into pCHF3-GFP digested with *EcoRI* and *Sac*I to generate pCHF3-V3-GFP (pV3-GFP) using In-Fusion Cloning according to the manufacturer’s instructions. The full-length V3 ORF was inserted into the PVX vector digested with *ClaI* and *SalI* to generate PVX-V3. The pCHF3-35S-GFP, pCHF3-p19, PVX-βC1, and a PVX-based expression vector for PTGS suppression assays have been described previously (Li et al., 2015; Xiong et al., 2009), as has the PVX-based expression vector PVX-βC1 for TGS suppression assays (Yang et al., 2011). All primers used in this study can be found in Supplementary table 1.

### Sequence analysis

The ViralORFfinder platform was constructed by Shiny, an R package used to build interactive web applications (https://shiny.rstudio.com). The NCBI ORFfinder (https://www.ncbi.nlm.nih.gov/orffinder/) was used to identify ORFs for each uploaded virus, and the R package Gviz (Hahne and Ivanek, 2016) was used for visualization; for Supplementary figures 1 and 3, 1.2-mer genomic sequences were used. To investigate the conservation of a ORF-encoded protein of choice, this tool creates a small database with the translated sequences from the inputted viral sequences, and BLASTp is used to identify proteins with high identity (e-value ≤ 0.05). Phylogenetic trees were obtained by the R package DECIPHER and visualized by ggtree. Multiple sequence alignments were constructed by ClustalW and visualized by the R package ggmsa (https://cran.r-project.org/web/packages/ggmsa/vignettes/ggmsa.html). Names and NCBI accession numbers of virus species used in this work are listed in Supplementary tables 2-6.

### Prediction of domains or signals in protein sequences

The prediction of transmembrane domains (TM) was performed by TMHMM (http://www.cbs.dtu.dk/services/TMHMM/) and Phobius (https://phobius.sbc.su.se/). The prediction of nuclear localization signal (NSL) was performed by cNLS Mapper (http://nlsmapper.iab.keio.ac.jp/cgi-bin/NLS_Mapper_form.cgi; (Kosugi et al., 2009a; Kosugi et al., 2009b), The prediction of chloroplast transit peptide (cTP) was performed by ChloroP (http://www.cbs.dtu.dk/services/ChloroP/).

### Confocal microscopy

Confocal microscopy was performed using a Leica TCS SP8 point scanning confocal microscope (Figure 2, 3h upper panel, Supplementary figures 8, 9, 10a, and 10d) or Zeiss LSM980 confocal microscope (Carl Zeiss) (Figure 3e, 3h lower panel, Supplementary figure 10b, c), with the preset settings for GFP (Ex: 488 nm, Em: 500-550 nm), RFP (Ex: 561 nm, Em: 570-620 nm), YFP (Ex:514 nm, Em: 515–570 nm), or mCherry (Ex: 594 nm, Em: 597–640 nm). For co-localization imaging, the sequential scanning mode was used.

### Mitochondrial staining

To visualize mitochondria, staining with 250 nM MitoTracker^®^ Red CMXRos (Invitrogen) was used. The chemical was infiltrated 10-30 min before imaging. The stock solution (1 mM) was prepared by dissolving the corresponding amount of MitoTracker^®^ in dimethylsulfoxide (DMSO). The working solution was prepared by diluting the stock solution in water or infiltration buffer (10 mM MgCl2, 10 mM MES (pH 5.6), and 100 μM acetosyringone). The MitoTracker^®^ red fluorescence was imaged using a Leica TCS SP8 point scanning confocal microscope with the following settings: Ex: 561 nm, Em: 570-620 nm.

### DNA and RNA extraction and qPCR/qRT-PCR

Total DNA was extracted from infected plants using the CTAB method. Total RNA was extracted from collected plant leaves using TaKaRa MiniBEST Universal RNA Extraction Kit (Takara, Japan). 1 ug of total RNA was reverse-transcribed into cDNA using PrimeScript^™^ RT reagent Kit with gDNA Eraser (Perfect Real Time) (Takara, Japan). qPCR or qRT-PCR was performed using TB Green^®^ Premix Ex Taq^™^ II (Takara, Japan). 25S RNA and *NbActin2* were used as internal references for DNA and RNA normalization, respectively.

### 5’ rapid amplification of cDNA ends (RACE)

Total RNA extracted from TYLCV (TYLCV-BJ, Accession No. MN432609)-infected *N. benthamiana* plants was used for 5’ RACE with SMARTer RACE 5’/3’ Kit (Takara, Japan) according to the manual booklet.

### 3,3’-diaminobenzidine (DAB) staining

For DAB staining, systemic leaves of infected plants were incubated in DAB solution (1 mg/mL, pH 3.8) for 10h at 25°C, then boiled for 5-10 min and decolorized in 95% ethanol.

### Protein extraction and western blotting

Total protein was isolated from infiltrated leaf patches with protein extraction buffer (containing 50 mM Tris-HCl (pH 6.8), 4.5% (m/v) SDS, 7.5% (v/v) 2-Mercaptoethanol, 9 M carbamide). Immunoblotting was performed with primary mouse polyclonal antibodies, followed by anti-mouse IgG HRP-linked antibodies (1:5000; Cell Signaling Technology, USA); primary antibodies used are as follows: anti-GFP (1:5000; ROCHE, USA), and custom-made anti-PVX CP (1:5000) and anti-TYLCV CP (1:5000) (Wu and Zhou, 2005; Wu et al., 2012).

## ACKNOWLEDGEMENTS

This work was supported by the National Natural Science Foundation of China (31930089), the Strategic Priority Research Program of the Chinese Academy of Sciences (Grant No. XDB27040206), and the Shanghai Center for Plant Stress Biology from the Chinese Academy of Sciences. The authors thank all members of Rosa Lozano-Duran’s lab and Alberto Macho’s lab for fruitful discussions, Xinyu Jian, Aurora Luque, and the PSC Cell Biology Facility for technical assistance, Alberto Macho for critical reading of the manuscript, and Dr. Michael M. Goodin (University of Kentucky, USA) and Prof. David Baulcombe (University of Cambridge) for kindly sharing materials.

## SUPPLEMENTARY MATERIAL

- Supplementary figures 1-12
- Supplementary tables 1-6

## Notes

### Competing Interest Statement

The authors have declared no competing interest.

