## Supplementary material for "Geminiviral genomes encode additional proteins with specific subcellular localizations and virulence function"

[1]State Key Laboratory for Biology of Plant Diseases and Insect Pests, Institute of Plant Protection, Chinese Academy of Agricultural Sciences, Beijing, 100193, China [2]Shanghai Center for Plant Stress Biology, CAS Center for Excellence in Molecular Plant Sciences, Chinese Academy of Sciences, Shanghai 201602, China. [3]University of the Chinese Academy of Sciences, Beijing 100049, China. [4]State Key Laboratory of Rice Biology, Institute of Biotechnology, Zhejiang University, Hangzhou, Zhejiang, 310058, China [5]Department of Plant Biochemistry, Centre for Plant Molecular Biology (ZMBP), Eberhard Karls University, D-72076 Tübingen, Germany.

*These authors contributed equally to this work.

This file contains:

- Supplementary figures 1-12
- Supplementary tables 1-6

### SUPPLEMENTARY FIGURES

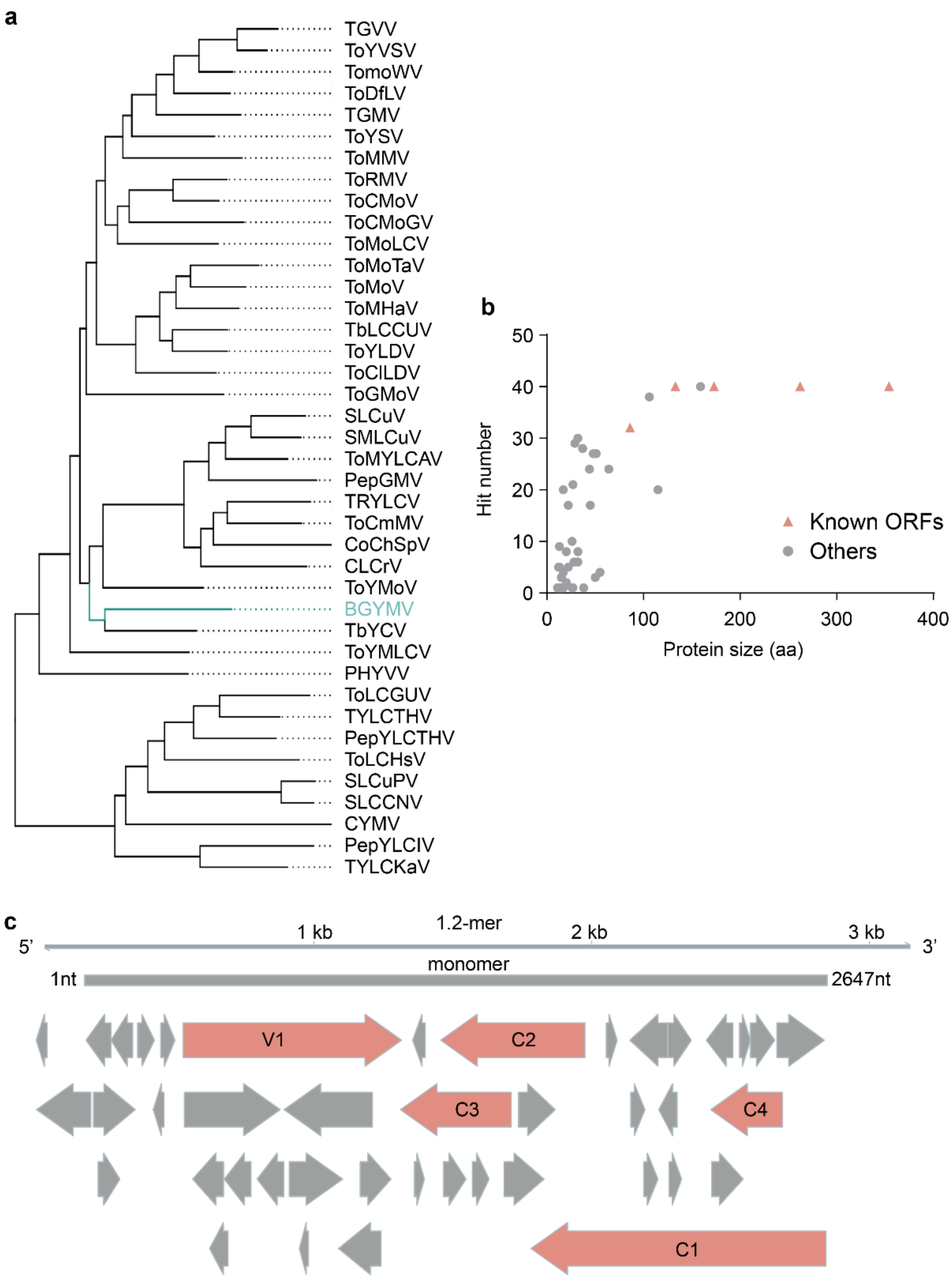

**Supplementary figure 1. Prediction of ORFs in the *Bean golden yellow mosaic virus* (BGYMV; gen. *Begomovirus*) genome.** a. Phylogenic tree based on the full genomic sequences of selected bipartite begomoviruses; for a full list, see Supplementary table 2. All sequences were downloaded from Genbank. BGYMV is indicated in blue. b. Schematic view of predicted ORFs (≥ 30 nt) in the BGYMV genome. A 1.2-mer was used for ORF prediction (see Methods section). c. Correlation between the size of the BGYMV ORFs-encoded proteins and their representation in the selected set of bipartite begomoviruses (a). In (b) and (c), arrows indicate ORFs; the known ORFs (C1, C2, C3, C4, and V1) are indicated in pink.

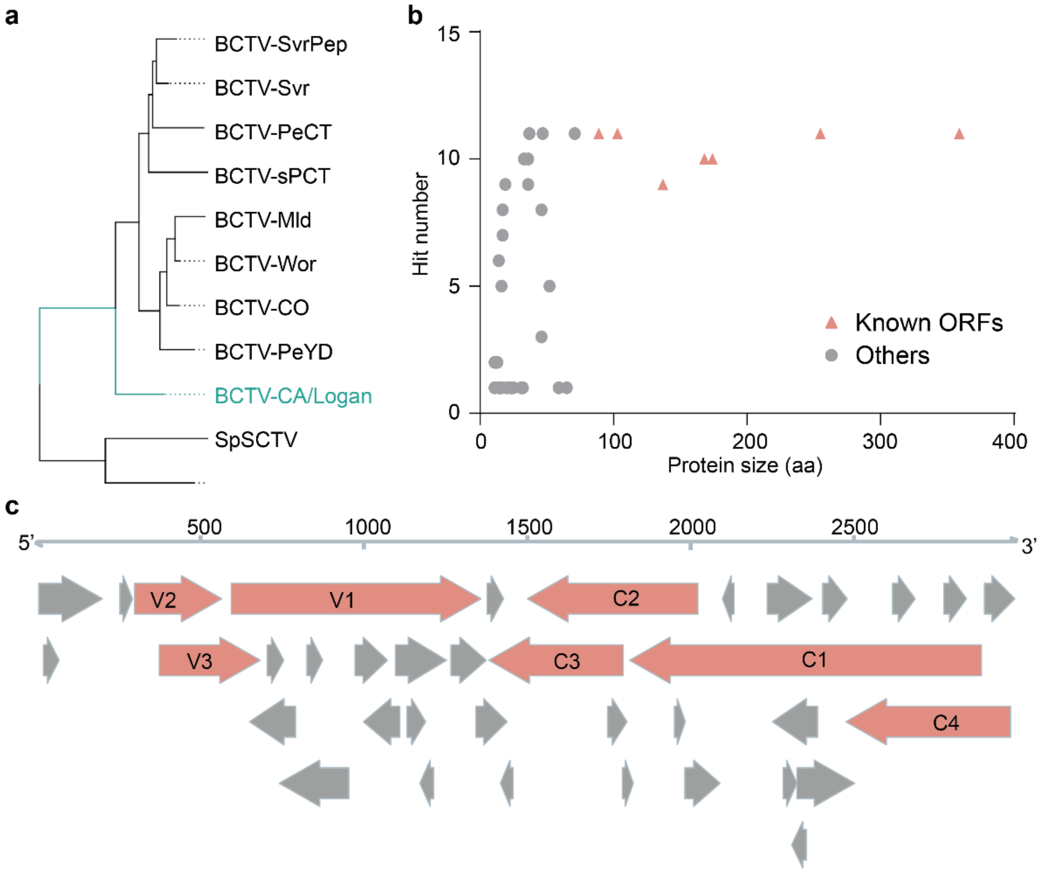


**Supplementary figure 2. Prediction of ORFs in the *Beet curly top virus* California/Logan strain (BCTV-AC/Logan; gen. *Curtovirus*) genome.** a. Phylogenic tree based on full genomic sequences of selected curtoviruses; for a full list, see Supplementary table 3. BCTV-CA/Logan is indicated in blue. b. Schematic view of predicted ORFs (≥ 30 nt) in BCTV-CA/Logan genome. c. Correlation between the size of the BCTV-CA/Logan ORFs-encoded proteins and their representation in the selected set of curtoviruses (s). In (b) and (c), arrows indicate ORFs; the known ORFs (C1, C2, C3, C4, V1, V2, and V3) are indicated in pink.

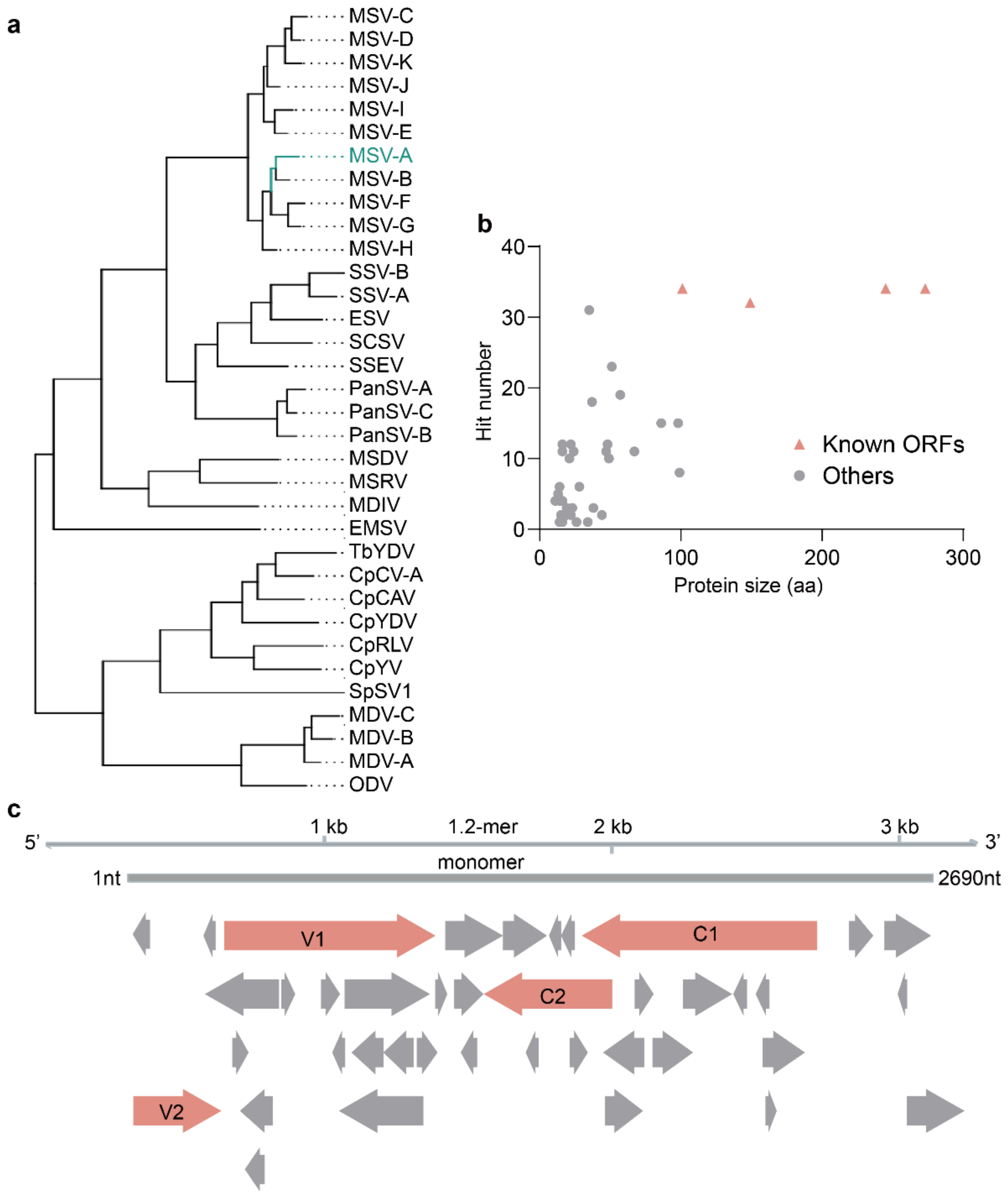


**Supplementary figure 3. Prediction of ORFs in the *Maize streak virus* A strain (MSV-A; gen. *Mastrevirus*) genome.** a. Phylogenic tree based on full genomic sequences of selected mastreviruses; for a full list, see Supplementary table 4. All sequences were downloaded from Genbank. MSV is indicated in blue. b. Schematic view of predicted ORFs (≥ 30 nt) in the MSV-A genome. A 1.2-mer was used for ORF prediction (see Methods section). c. Correlation between the size of the MSV-A ORFs-encoded proteins and their representation in the selected set mastreviruses (a). In (b) and (c), arrows indicate ORFs; the known ORFs (C1, C2, V1, and V2) are indicated in pink.

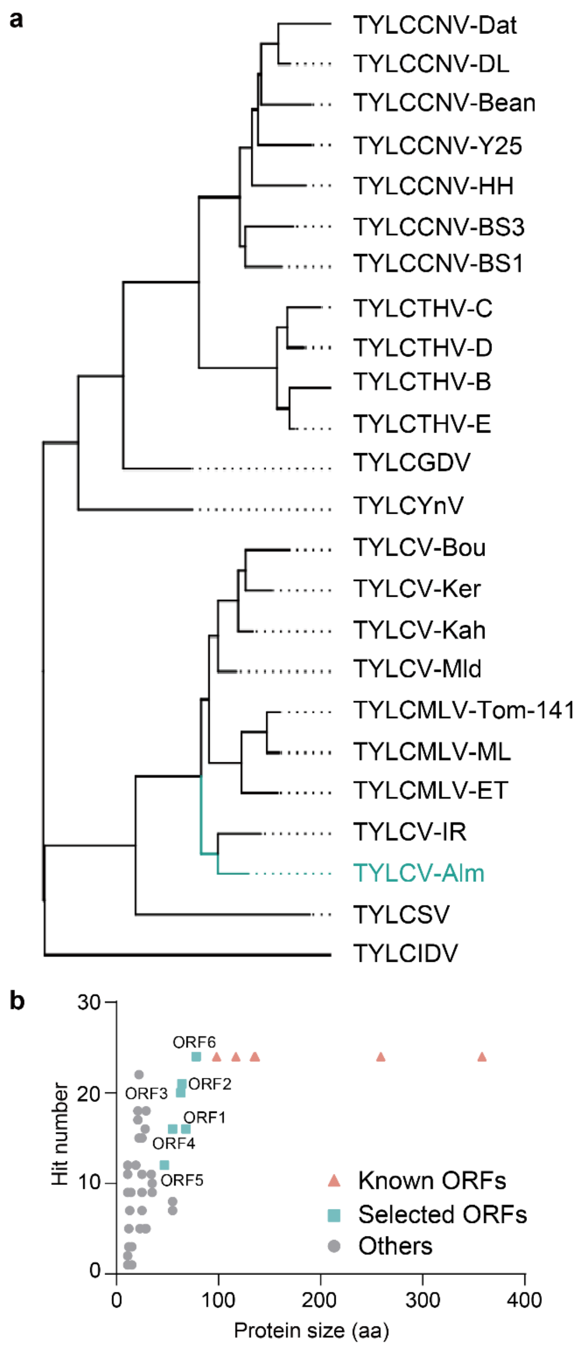


**Supplementary figure 4. Prediction of ORFs in the *Tomato yellow leaf curl virus* (TYLCV-Alm; gen. *Begomovirus*) genome.** a. Phylogenic tree based on full genomic sequences of selected monopartite tomato-infecting begomoviruses; for a full list, see Supplementary table 5. All sequences were downloaded from Genbank. TYLCV-Alm is indicated in blue. b. Correlation between the size of the TYLCV-Alm ORFs-encoded proteins and their representation in the selected set monopartite tomato-infecting begomoviruses (a). Known ORFs (C1, C2, C3, C4, V1 and V2) are indicated in pink; selected novel ORFs (ORF1-6) are indicated in blue.

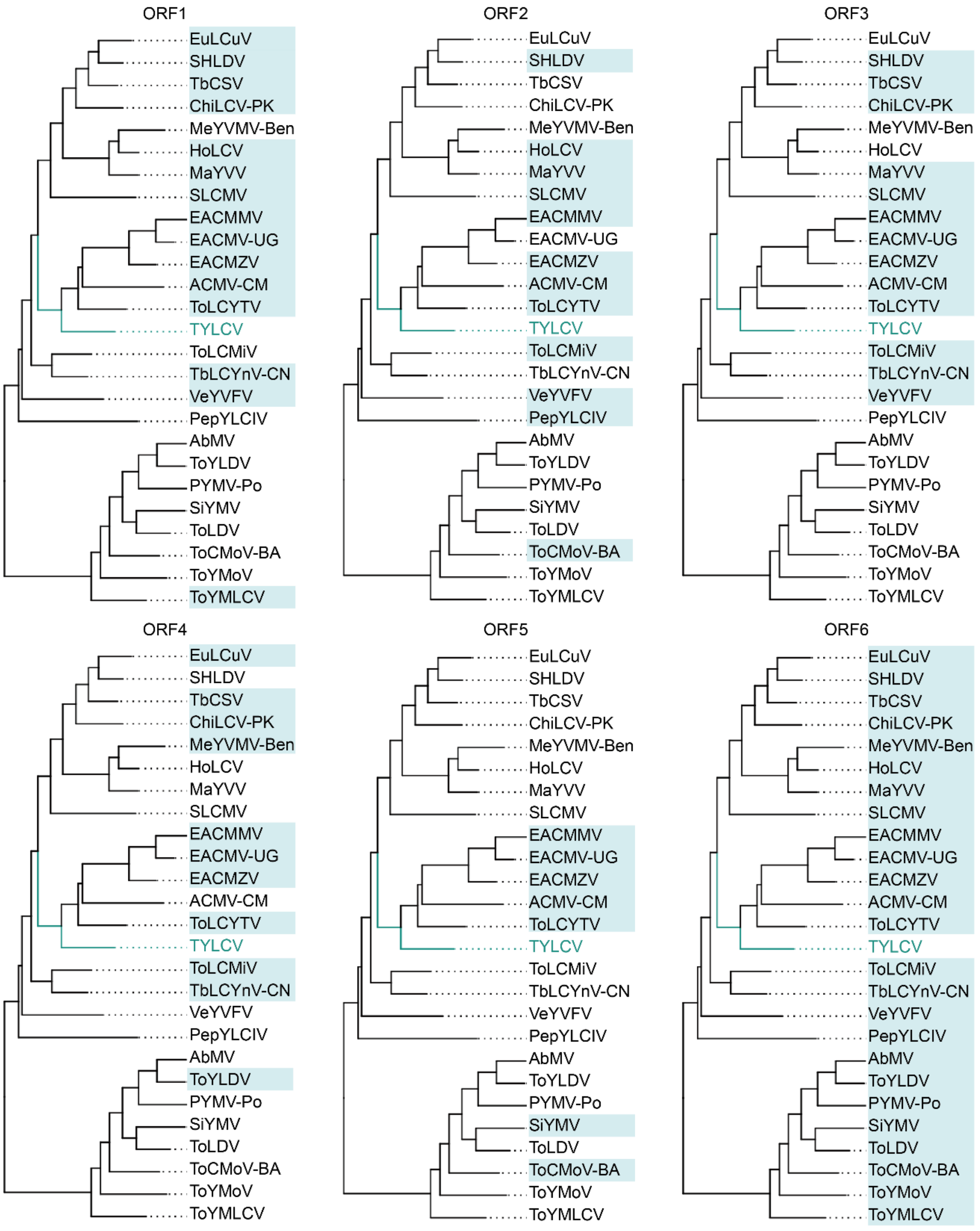


**Supplementary figure 5. Distribution of ORF1-6 from TYLCV in the selected subset of mono- and bipartite begomoviruses.** Blue boxes indicate presence of a given ORF (ORF1-6) described in TYLCV (see Supplementary figure 4). For a full list of species, see Supplementary table 6. All sequences were downloaded from Genbank.

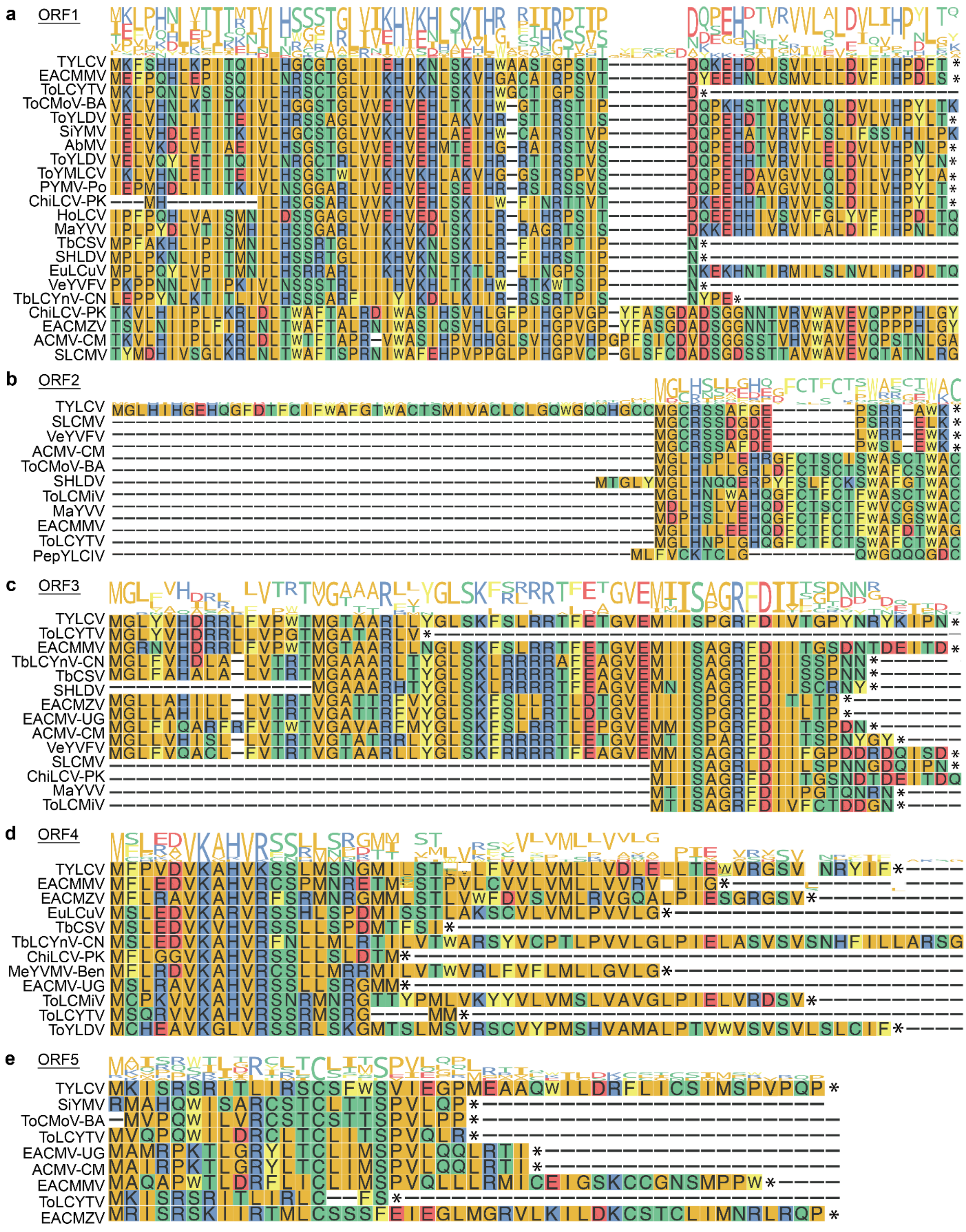
a
ORF1
b
ORF2
c
ORF3
d
ORF4
e
ORF5

**Supplementary figure 6. Multiple sequence alignment of the proteins encoded by ORF1-5 in different begomoviruses.** Multiple sequence alignment of the proteins encoded by ORF1 (a), ORF2 (b), ORF3 (c), ORF4 (d), and ORF5 (e). Multiple sequence alignments were performed by ClustalW. Asterisks indicate stop codons.

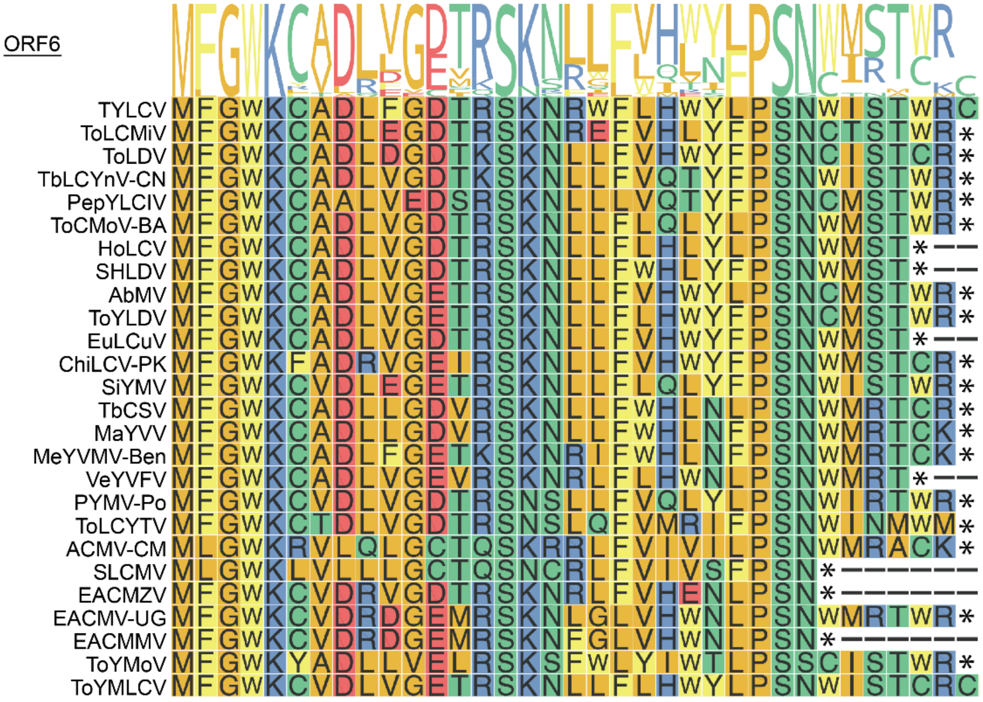


**Supplementary figure 7. Multiple sequence alignment of the protein encoded by ORF6 in different begomoviruses.** Multiple sequence alignments were performed by ClustalW. Asterisks indicate stop codons.

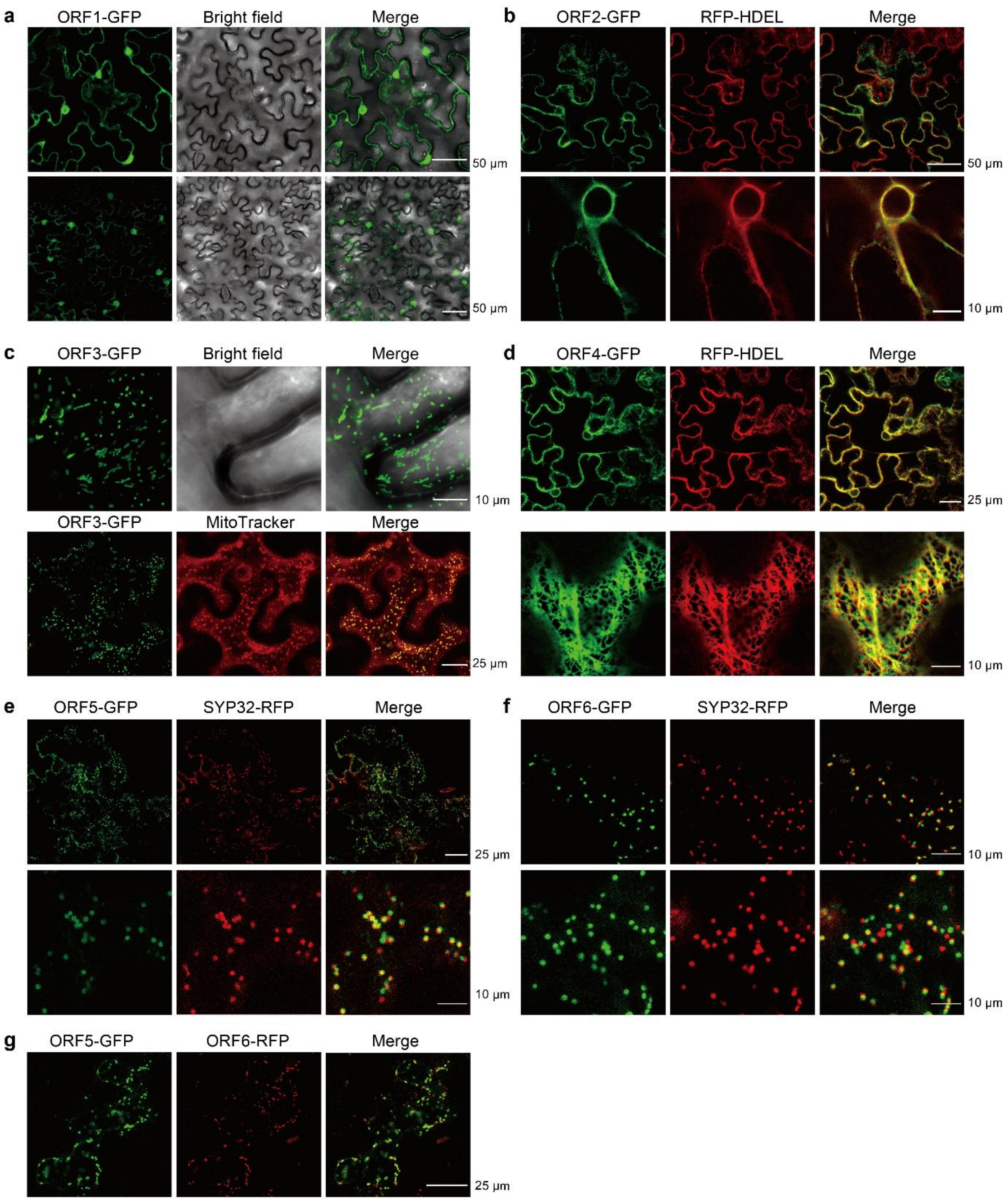


**Supplementary figure 8. Additional confocal images showing the subcellular localization of GFP-tagged proteins encoded by the novel ORFs from TYLCV (ORF1-6) transiently expressed in *N. benthamiana* leaves (related to Figure 2).** RFP-HDEL: ER marker; SYP32-RFP: cis-Golgi marker; MitoTracker Red: mitochondrial staining. The size of scale bars is indicated. These experiments were repeated at least three times with similar results; representative images are shown. The TYLCV isolate used in these experiments is TYLCV-Alm.

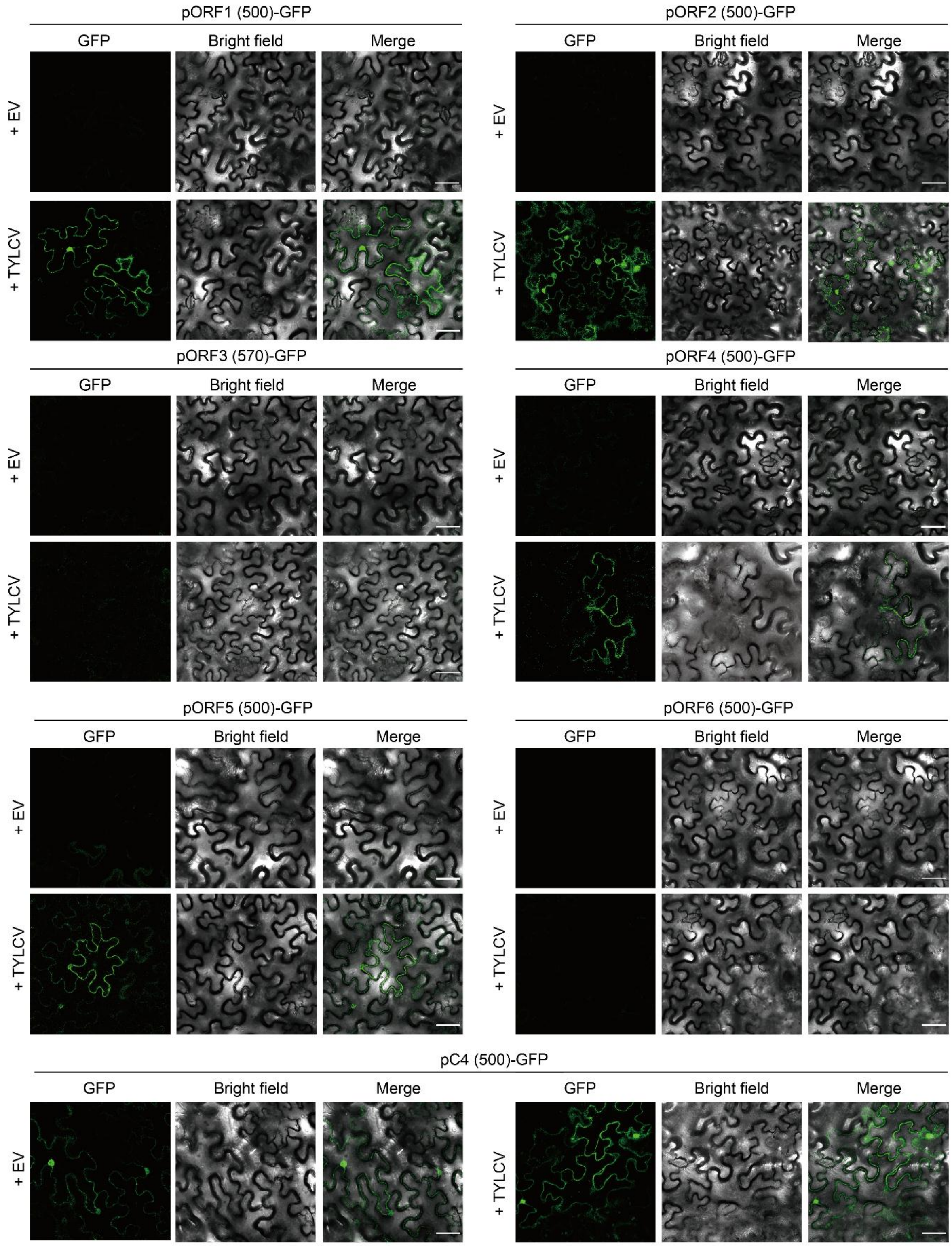
pORF1 (500)-GFP
pORF2 (500)-GFP
GFP
Bright field
Merge
+ EV
+ TYLCV
pORF3 (570)-GFP
pORF4 (500)-GFP
pORF5 (500)-GFP
pORF6 (500)-GFP
pC4 (500)-GFP

**Supplementary figure 9. Promoter activity assay of the sequences upstream of ORF1-6.** *A. tumefaciens* clones harboring pORF-GFP constructs were used to transiently transform *N. benthamiana* leaves in the presence (+TYLCV) or absence (+EV) of TYLCV. EV: empty vector. Scale bar: 25 µm. The numbers in brackets indicate the length of the sequenced used, upstream of the ATG of each ORFs, in nt. These experiments were repeated three times with similar results; representative images are shown. The TYLCV isolate used in these experiments is TYLCV-Alm.

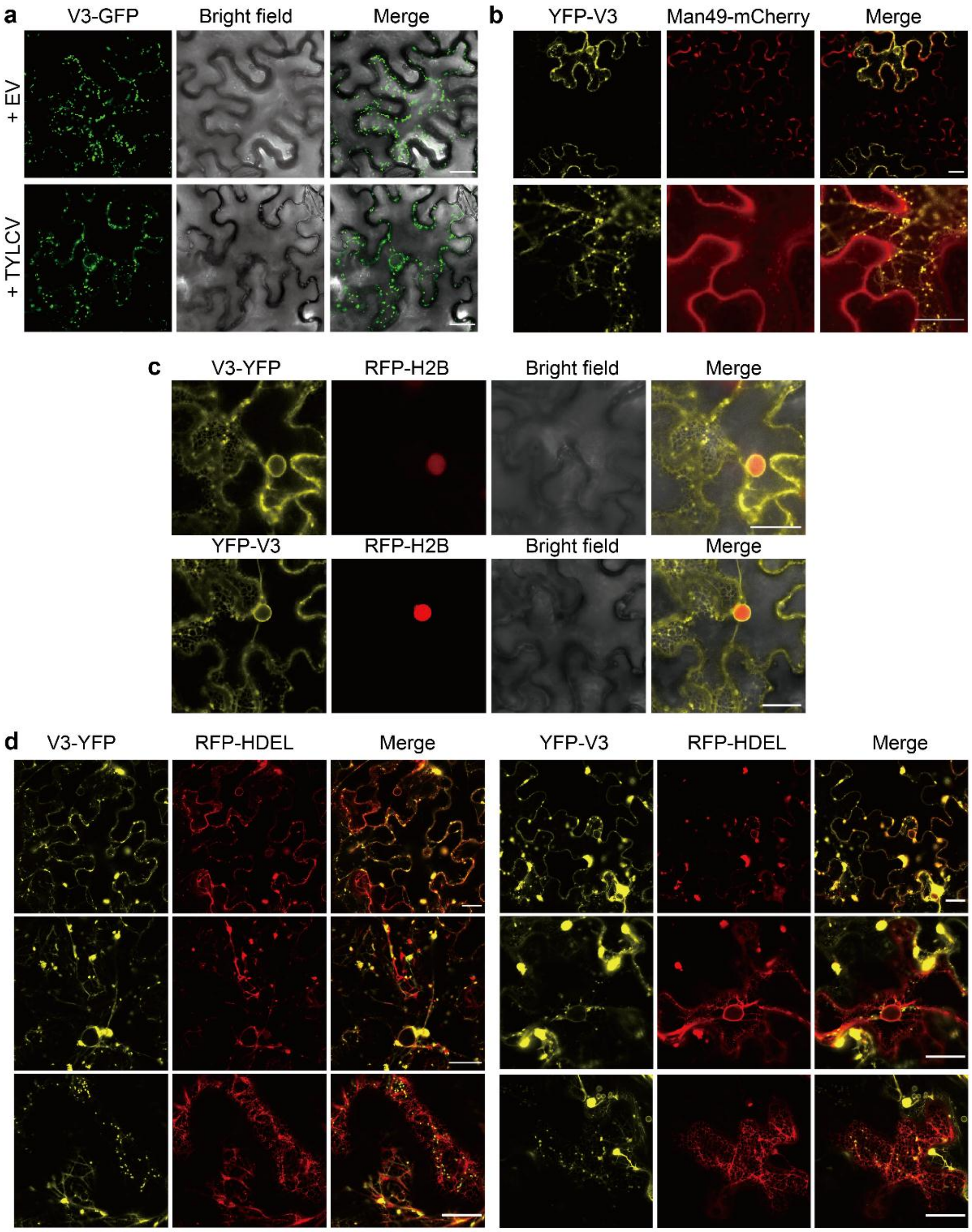


**Supplementary figure 10. Additional confocal images showing the subcellular localization of V3 with fluorescent protein tags at its N- or C-terminus.** a. Subcellular localization of V3-GFP in the presence (+TYLCV) or absence (+EV) of TYLCV. EV: empty vector. Scale bar: 25 μm. b. Co-localization of YFP-V3 with the cis-Golgi marker Man49-mCherry. Scale bar: 20 μm. c. Subcellular localization of V3-YFP or YFP-V3 transiently expressed in RFP-H2B transgenic *N. benthamiana* leaves. Scale bar: 20 μm. d. Co-localization of V3-YFP or YFP-V3 with the ER marker RFP-HDEL. Scale bar: 25 μm. a-d. These experiments were repeated at least three times with similar results; representative images are shown.

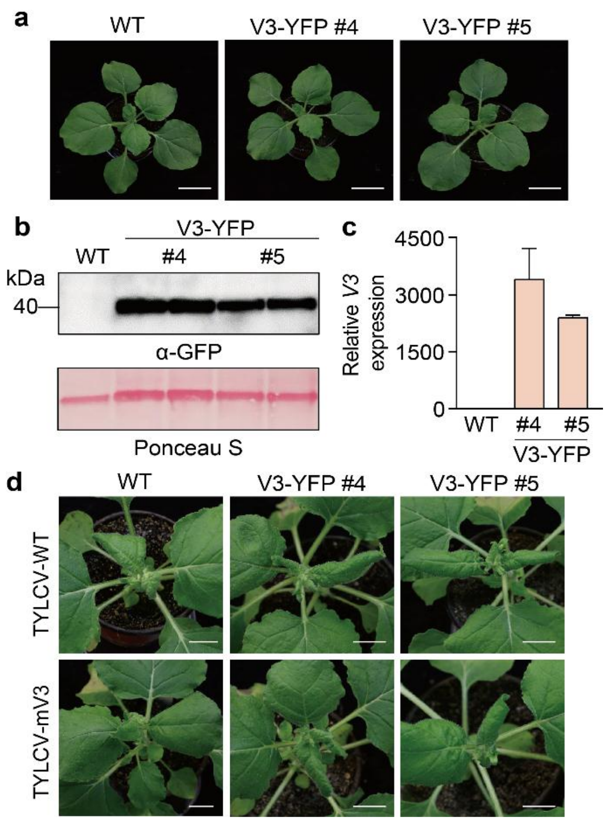


**Supplementary figure 11. Characterization of transgenic *N. benthamiana* lines expressing V3.** a. Phenotype of 5-week-old p35S:*V3-YFP* T1 transgenic *N. benthamiana* lines (#4 and #5) compared to the wild type (WT). Bar= 20 µm. b. Western blot showing YFP protein accumulation in the plants in (a). The corresponding Ponceau S staining of the large RuBisCO subunit serves as loading control. c. Relative V3 expression levels in WT and V3-YFP transgenic lines measured by qRT-PCR. *NbActin2* was used as internal reference. d. Symptoms of V3-YFP transgenic *N. benthamiana* plants infected with TYLCV-WT or TYLCV-mV3 at 11 dpi.

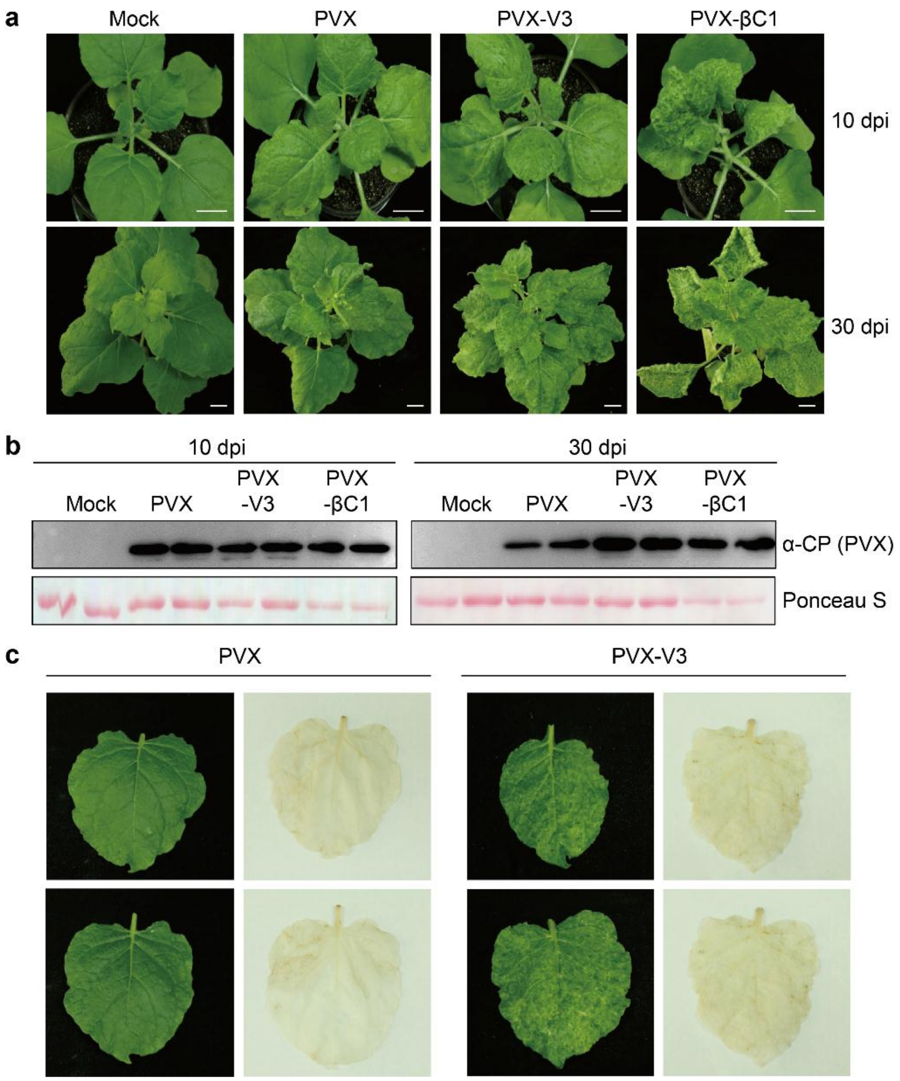


**Supplementary figure 12. V3 enhances the pathogenicity of PVX.** a. Symptoms of *N. benthamiana* plants infected with PVX, PVX-V3, PVX-βC1 (as positive control), or mock-inoculated at 10 dpi (upper panel) or 30 dpi (lower panel). b. Western blot showing the accumulation of PVX CP in systemic leaves from (a). The corresponding Ponceau S staining of the large RuBisCO subunit serves as loading control. c. DAB staining of *N. benthamiana* leaves infiltrated with PVX or PVX-V3.

### SUPPLEMENTARY TABLES

#### Supplementary table 1. Primers used in this work.

| Primer name | Sequence (5'-3') | Purpose |
|---|---|---|
| ORF1-F | CACCATGGGCCTTCACATCCACGG | To generate TOPO-ORF1-NS |
| ORF1-NS-R | TTTCCACGCCCGTCTCGAAG | To generate TOPO-ORF1-NS |
| ORF2-F | CACCATGAAATTTTCTCATCACTTGAAAC | To generate TOPO-ORF2-NS |
| ORF2-NS-R | GGTAAAGTCTGGATGGATGAA | To generate TOPO-ORF2-NS |
| ORF3-F | GGGGACAAGTTTGTACAAAAAAGCAGGCTTCATGGGCCTGTACGTCCATGA | To generate pDONR-ORF3-NS |
| ORF3-NS-R | GGGGACCACTTTGTACAAGAAAGCTGGGTCATTAGGGATCTTATATCTG | To generate pDONR-ORF3-NS |
| ORF4-F | CACCATGTTCCCCGTGGATGTGAAG | To generate TOPO-ORF4-NS |
| ORF4-NS-R | AAAAATATATCGATTTAACAC | To generate TOPO-ORF4-NS |
| ORF5-F | CACCATGAAAATATCAAGAAGCAGA | To generate TOPO-ORF5-NS |
| ORF5-NS-R | CGGTTGCGGTACTGGGCTCAT | To generate TOPO-ORF5-NS |
| ORF6-F | CACCATGTTCGGATGGAAATGTGCTG | To generate TOPO-ORF6-NS |
| ORF6-NS-R | TTTCCTAACATATCCCAATTGT | To generate TOPO-ORF6-NS |
| SYP32-F | CACCATGTCGGCAAGGCATGGGCAAT | To generate TOPO-SYP32-NS |
| SYP32-NS-R | TGCCACGAAGAAGAGGAAAATCAT | To generate TOPO-SYP32-NS |
| ORF1pro500-F | GGGGACAAGTTTGTACAAAAAAGCAGGCTTCAGAGGCATGCGTACATGCCA | To generate pDONR-pORF1(500) |
| ORF1pro500-R | GGGGACCACTTTGTACAAGAAAGCTGGGTCGTAAAGTCCAGTCTTATGAG | To generate pDONR-pORF1(500) |
| ORF2pro500-F | GGGGACAAGTTTGTACAAAAAAGCAGGCTTCAACCACGACATCATTTCCATTC | To generate pDONR-pORF2(500) |
| ORF2pro500-R | GGGGACCACTTTGTACAAGAAAGCTGGGTCGCAACAGTTATTGGTGGGC | To generate pDONR-pORF2(500) |
| ORF3pro570-F | GGGGACAAGTTTGTACAAAAAAGCAGGCTTCACTGGATTAGAGGCATGCGT | To generate pDONR-pORF3(570) |
| ORF3pro570-R | GGGGACCACTTTGTACAAGAAAGCTGGGTCCGAAAGCCCAGAATATACAGAATG | To generate pDONR-pORF3(570) |
| ORF4pro500-F | GGGGACAAGTTTGTACAAAAAAGCAGGCTTCATTACCGGATGGCCGCGCCT | To generate pDONR-pORF4(500) |
| ORF4pro500-R | GGGGACCACTTTGTACAAGAAAGCTGGGTCCAGGGCTTCGATACATTCTG | To generate pDONR-pORF4(500) |
| ORF5pro500-F | GGGGACAAGTTTGTACAAAAAAGCAGGCTTCGAATCTGTTCACGGATTTCG | To generate pDONR-pORF5(500) |
| OR5pro500-R | GGGGACCACTTTGTACAAGAAAGCTGGGTCCCATCCAGACTTTACCTAA | To generate pDONR-pORF5(500) |
| ORF6pro500-F | GGGGACAAGTTTGTACAAAAAAGCAGGCTTCTGTACCACGCATCATTACTG | To generate pDONR-pORF6(500) |
| ORF6pro500-R | GGGGACCACTTTGTACAAGAAAGCTGGGTCTCAGGCAGCTAAGAGCTCAAC | To generate pDONR-pORF6(500) |

| | | |
|---|---|---|
| C4pro500-F | GGGGACAAGTTTGTACAAAAAAGCAGGCTTCAGATTCAGGAAATTC ATTTAG | To generate pDONR-pC4(500) |
| C4pro500-R | GGGGACCACTTTGTACAAGAAAGCTGGGTCTCTCGTGGAGTTCTCT GCAAAC | To generate pDONR-pC4(500) |
| V3-GSPs-R | GATTACGCCAAGCTTCTCTCTCTAAAGAGGAAGCACTTTCCC | For 5'RACE |
| V3-nGSPs-R | GATTACGCCAAGCTTGGGGAACCACATCTCCATGTGC | For 5'RACE |
| q-25S-rRNA-F | ATAACCGCATCAGGTCTCCA | For qPCR |
| q-25S-rRNA-R | CCGAAGTTACGGATCCATTT | For qPCR |
| q-TYLCV-F | CCCTCAAAGCTCTATGGCAATCGG | For qPCR |
| q-TYLCV-R | CAGTGACGTCTGTGGAACCCTC | For qPCR |
| qV3-F | ATGTTCGGATGGAAATGTGCTGA | For qPCR |
| qV3-R | GTTTGCAGAGAACTCCACGAGAA | For qPCR and reverse transcription |
| NbActin2-qPCR-F | AAAGACCAGCTCATCCGTGGAGAA | For qPCR |
| NbActin2-qPCR-R | TGTGGTTTCATGAATGCCAGCAGC | For qPCR |
| IF V3pro833 GFP-F | GCTATGACATGATTACGAATTCGCAGCCACAGTCTAGGTC | To generate pCHF3-V3-GFP vector |
| IF V3pro833 GFP-R | GGATCCCCGGGTACCGAGCTCTCAGGCAGCTAAGAGCTC | To generate pCHF3-V3-GFP vector |
| IF V3pro833 GUS-F | CCAAGCTGGCGCGCCAAGCTTGCAGCCACAGTCTAGGTC | To generate pINT121-V3-GUS vector |
| IF V3pro833 GUS-R | GACTGACCTACCCGGGGATCCTCAGGCAGCTAAGAGCTCAAC | To generate pINT121-V3-GUS vector |
| mV3-F | TTAGCTGCCTGAACGTTCGGATGGAAATG | To generate TYLCV-mV3 |
| mV3-R | CCATCCGAACGTTCAGGCAGCTAAGAGCTCA | To generate TYLCV-mV3 |
| IF-PVX(ClaI)-V3-F | GCACCAGCTAGCATCGAT ATGTTCGGATGGAAATGTG | To generate PVX-V3 |
| IF-PVX(SalI)-V3-R | GTTCATCGGCGGTCGAC TTATTTCCTAACATATCCCAATTG | To generate PVX-V3 |
| 221-V3-F | GGGGACAAGTTTGTACAAAAAAGCAGGCTTCATGTTCGGATGGAAA TG | To generate 221-V3 |
| 221-V3-R | GGGGACCACTTTGTACAAGAAAGCTGGGTCTTTCCTAACATATCCCA ATTG | To generate 221-V3 |

### Supplementary table 2. Selected subset of bipartite begomoviruses (from Supplementary figure 1).

| Species | Isolate | Accession number | Virus Abbrev. |
|---|---|---|---|
| *Squash leaf curl Philippines virus* | Philippines/Munoz | DNA-A: AB085793; DNA-B: AB085794 | SLCuPV |
| *Pepper yellow leaf curl Indonesia virus* | Indonesia/2005 | DNA-A: AB267834; DNA-B: AB267835 | PepYLCIV |

| | | | |
|---|---|---|---|
| *Tomato mottle Taino virus* | Cuba | DNA-A: AF012300; DNA-B: AF012301 | ToMoTaV |
| *Tomato yellow leaf curl Thailand virus* | Thailand/2/A | DNA-A: AF141922; DNA-B: AF141897 | TYLCTHV/A |
| *Tomato rugose mosaic virus* | Brazil/Uberlandia 1/1996 | DNA-A: AF291705; DNA-B: AF291706 | ToRMV |
| *Squash mild leaf curl virus* | United States/Imperial Valley/1979 | DNA-A: AF421552; DNA-B: AF421553 | SMLCuV |
| *Cotton leaf crumple virus* | Mexico/Sonora/1991/Arizona | DNA-A: AF480940; DNA-B: AF480941 | CLCrV/AZ |
| *Tomato chlorotic mottle virus* | Brazil/Seabra 1/1996/Bahia | DNA-A: AF490004; DNA-B: AF491306 | ToCMoV/BA |
| *Squash leaf curl China virus* | Vietnam/B/China | DNA-A: AF509743; DNA-B: AF509742 | SLCCNV/CN |
| *Tomato yellow leaf curl Kanchanaburi virus* | Thailand/Kanchanaburi 1/2001 | DNA-A: AF511529; DNA-B: AF511528 | TYLCKaV |
| *Tomato leaf curl Gujarat virus* | India/Varanasi/2001 | DNA-A: AY190290; DNA-B: AY190291 | ToLCGUV |
| *Tomato yellow margin leaf curl virus* | Venezuela/Merida/57 | DNA-A: AY508993; DNA-B: AY508994 | ToYMLCV |
| *Tomato mild yellow leaf curl Aragua virus* | Venezuela/10/2003 | DNA-A: AY927277; DNA-B: EF547938 | ToMYLCAV |
| *Tomato yellow spot virus* | Brazil/Bicas 2/1999 | DNA-A: DQ336350; DNA-B: DQ336351 | ToYSV |
| *Tomato golden mottle virus* | Mexico/San Luiz Potosi/2005 | DNA-A: DQ520943; DNA-B: DQ406674 | ToGMoV |
| *Tomato yellow vein streak virus* | Brazil/Potato/1983 | DNA-A: EF417915; DNA-B: EF417916 | ToYVSV |
| *Tomato leaf curl Hsinchu virus* | China/Hainan/Ramie/2007 | DNA-A: EU596959; DNA-B: EU596960 | ToLCHsV |
| *Tomato mild mosaic virus* | Brazil/Paty do Alferes 58/2005 | DNA-A: EU710752; DNA-B: EU710753 | ToMMV |
| *Tomato common mosaic virus* | Brazil/Coimbra 22/2007 | DNA-A: EU710754; DNA-B: EU710755 | ToCmMV |
| *Tomato yellow leaf distortion virus* | Cuba/5E17/2007 | DNA-A: FJ174698; DNA-B: FJ999999 | ToYLDV |
| *Tobacco yellow crinkle virus* | Cuba/2007 | DNA-A: FJ213931; DNA-B: HQ896204 | TbYCV |
| *Tomato chlorotic leaf distortion virus* | Venezuela/Zulia/2004 | DNA-A: HQ201952; DNA-B: HQ201953 | ToClLDV |
| *Tomato golden vein virus* | Brazil/Ita1220/2003 | DNA-A: JF803254; DNA-B: JF803265 | TGVV |
| *Tomato rugose yellow leaf curl virus* | Uruguay/Salto Grande/U2/2009 | DNA-A: JN381819; DNA-B: JN381814 | TRYLCV |
| *Tomato dwarf leaf virus* | Argentina/Pichanal 397/2008 | DNA-A: JN564749; DNA-B: JN564750 | ToDfLV |
| *Tomato mottle wrinkle virus* | Argentina-Pichanal_400-2008 | DNA-A: JQ714137; DNA-B: JQ714138 | ToMoWV |
| *Tomato golden mosaic virus* | Brazil/Common/1984 | DNA-A: K02029; DNA-B: K02030 | TGMV |

| | | | |
|---|---|---|---|
| *Tomato yellow mottle virus* | Costa Rica/2003 | DNA-A: KC176780; DNA-B: KC176781 | ToYMoV |
| *Tomato mottle leaf curl virus* | Brazil/Jaiba 13/2008 | DNA-A: KC706615; DNA-B: JF803264 | ToMoLCV |
| *Cotton chlorotic spot virus* | Brazil/CampinaGrandeB012/2009 | DNA-A: KF358470; DNA-B: KF358471 | CoChSpV |
| *Tomato chlorotic mottle Guyane virus* | French Guyana-Mon2-GF455-2009 | DNA-A: KR263181; DNA-B: KR263172 | ToCMoGV |
| *Cotton yellow mosaic virus* | Benin-Gos_San2-2014 | DNA-A: KU683748; DNA-B: KU683750 | CYMV |
| *Tobacco leaf curl Cuba virus* | Cuba/VC/2015 | DNA-A: KX011471; DNA-B: KX011472 | TbLCCUV |
| *Pepper yellow leaf curl Thailand virus* | Thailand-WF_SPN_Pep-2015 | DNA-A: KX943290; DNA-B: KX943291 | PepYLCTHV |
| *Tomato mottle virus* | United States/Florida/1989 | DNA-A: L14460; DNA-B: L14461 | ToMoV |
| *Squash leaf curl virus* | United States/Imperial Valley/1979 | DNA-A: M38183; DNA-B: M38182 | SLCuV |
| *Pepper golden mosaic virus* | Mexico/Tamaulipas/Tamaulipas | DNA-A: U57457; DNA-B: AF499442 | PepGMV/Tam |
| *Pepper huasteco yellow vein virus* | Mexico/Tamaulipas | DNA-A: X70418; DNA-B: X70419 | PHYVV |
| *Tomato mosaic Havana virus* | Cuba/Quivican | DNA-A: Y14874; DNA-B: Y14875 | ToMHaV |
| *Bean golden yellow mosaic virus* | Dominican Republic/1987 | DNA-A: L01635; DNA-B: L01636 | BGYMV |

### Supplementary table 3. Selected subset of curtoviruses (from Supplementary figure 2).

| **Species** | **Isolate** | **Accession number** | **Virus Abbrev.** |
|---|---|---|---|
| *Beet curly top virus* | United States/CA/Logan/1985/California Logan | M24597 | BCTV/CA/Logan |
| *Beet curly top virus* | Mexico/Mild/2006/Mild | EU193175 | BCTV/Mld |
| *Beet curly top virus* | United States/Severe/Cfh/Severe | U02311 | BCTV/Svr |
| *Beet curly top virus* | United States/New Mexico/Severe/Pepper/2001/Severe pepper | FJ545686 | BCTV/SvrPep |
| *Beet curly top virus* | United States/Colorado/1995/Colorado | JN817383 | BCTV/CO |
| *Beet curly top virus* | United States/Mild/Worland/Worland | U56975 | BCTV/Wor |
| *Beet curly top virus* | United States/Spinach 3/1996/Spinach curly top | AY548948 | BCTV/ SpCT |
| *Beet curly top virus* | United States/New Mexico/Pepper/2005/Pepper curly top | EF501977 | BCTV/ PeCT |
| *Beet curly top virus* | United States/New Mexico/Pepper/2007/Pepper yellow dwarf | EU921828 | BCTV/ PeYD |
| *Horseradish curly top virus* | United States/Salinas/1988 | U49907 | HCTV |
| *Spinach severe curly top virus* | United States/Arizona/Spinach 0910/2009 | GU734126 | SpSCTV |

### Supplementary table 4. Selected subset of mastreviruses (from Supplementary figure 3).

| Species | Isolate | Accession number | Virus Abbrev. |
|---|---|---|---|
| *Chickpea chlorosis Australia virus* | Australia/2614/2010] | JN989422 | CpCAV |
| *Chickpea chlorosis virus* | Australia/3455C/2002/A | GU256530 | CpCV/A |
| *Chickpea redleaf virus* | Australia/Queensland 22/2003 | GU256532 | CpRLV |
| *Chickpea yellow dwarf virus* | Pakistan/PK37/ | KM377674 | CpYDV |
| *Chickpea yellows virus* | Australia/3489B/2002 | JN989439 | CpYV |
| *Eragrostis minor streak virus* | Namibia/Caprivi/g450/2009 | JF508490 | EMSV |
| *Eragrostis streak virus* | Zimbabwe/Guruwe 186/2007 | EU244915 | ESV |
| *Maize streak dwarfing virus* | ET-Adama Zuria-MV1-16 | MK329300 | MSDV |
| *Maize streak Reunion virus* | La Reunion/St Pierre/PR52/2009 | JQ624879 | MSRV |
| *Maize streak virus* | South Africa/A | Y00514 | MSV/A |
| *Maize streak virus* | South Africa/Worester/Plaas Staal B/g27b/2006/B | EU628597 | MSV/B |
| *Maize streak virus* | South Africa/Mt Edgecomb/Setaria/1988/C | AF007881 | MSV/C |
| *Maize streak virus* | South Africa/Rawsonville/1998/D | AF329889 | MSV/D |
| *Maize streak virus* | South Africa/Mitchelle Park/Natal A/g125/Dig/2006/E | EU628626 | MSV/E |
| *Maize streak virus* | Nigeria/IITA B/g88/Urochloa/2007/F | EU628629 | MSV/F |
| *Maize streak virus* | Mali/Mic25/Digitaria/1987/G | EU628631 | MSV/G |
| *Maize streak virus* | Nigeria/Lagbaka/g79/Setaria/2007/H | EU628638 | MSV/H |
| *Maize streak virus* | South Africa/New Germany/Natal/A/g217/Digitaria/2007/I | EU628639 | MSV/I |
| *Maize streak virus* | Zimbabwe/Mic24K/Pennisetum/1987/J | EU628641 | MSV/J |
| *Maize streak virus* | Uganda/Busia 4/Eustachys/2005/K | EU628643 | MSV/K |
| *Oat dwarf virus* | Germany/Saxena 25/2006 | AM296025 | ODV |
| *Panicum streak virus* | South Africa/Karino/1989/A | L39638 | PanSV/A |
| *Panicum streak virus* | Kenya/1990/B | X60168 | PanSV/B |
| *Panicum streak virus* | Zimbabwe/Guruwe 169/Urochloa/2006/C | EU224264 | PanSV/C |
| *Sugarcane chlorotic streak virus* | Nigeria/Sc-10/SR/1/2015 | KX787914 | SCSV |
| *Sugarcane streak Egypt virus* | Egypt/Aswan | AF037752 | SSEV |
| *Sugarcane streak virus* | South Africa/Natal/A | M82918 | SSV/A |
| *Sugarcane streak virus* | La Reunion/St Pierre/Pie/R5/Cenchrus/2006/B | EU244914 | SSV/B |
| *Sweet potato symptomless virus 1* | Kenya/Q44429/2005 | KY565231 | SpSV/1 |
| *Tobacco yellow dwarf virus* | Australia/1992 | M81103 | TbYDV |
| *Wheat dwarf India virus* | India/2010 | JQ361910 | WDIV |
| *Wheat dwarf virus* | Turkey/barley/A | AJ783960 | WDV/A |
| *Wheat dwarf virus* | Iran/2008/B | FJ620684 | WDV/B |
| *Wheat dwarf virus* | Hungary/Kompolt10/1/2010/C | JQ647455 | WDV/C |

### Supplementary table 5. Selected subset of monopartite begomoviruses causing tomato yellow leaf curl disease (from Supplementary figure 4).

| Species | Isolate | Accession number | Virus Abbrev. |
|---|---|---|---|

| | | | |
|---|---|---|---|
| *Tomato yellow leaf curl virus* | Spain/Almeria/Pepper/1999 | AJ489258 | TYLCV |
| *Tomato yellow leaf curl virus* | Iran/Genaveh 29/2006/Boushehr | GU076454 | TYLCV/Bou |
| *Tomato yellow leaf curl virus* | Iran/Iranshahr/1998/Iran | AJ132711 | TYLCV/IR |
| *Tomato yellow leaf curl virus* | Iran/Kahnooj/2007/Kahnoo | EU635776 | TYLCV/Kah |
| *Tomato yellow leaf curl virus* | Iran/Hormozgan 32/2006/Kerman | GU076442 | TYLCV/Ker |
| *Tomato yellow leaf curl virus* | Israel/1993/Mild | X76319 | TYLCV/Mld |
| *Tomato yellow leaf curl China virus* | China/Guangxi/Honghe | DNA-A: AF311734 | TYLCCNV/HH |
| *Tomato yellow leaf curl China virus* | China/Yunnan 10/Tobacco/2000/Baoshan | AJ319675 | TYLCCNV/BS1 |
| *Tomato yellow leaf curl China virus* | China/Yunnan 25/Tomato/2000 | DNA-A: AJ457985 | TYLCCNV |
| *Tomato yellow leaf curl China virus* | China/Yunnan 278/Malvastrum/2007/Baoshan3 | DNA-A: AM980509 | TYLCCNV/BS3 |
| *Tomato yellow leaf curl China virus* | China/Yunnan/Bean/2004/Bean | DQ256460 | TYLCCNV/Bea |
| *Tomato yellow leaf curl China virus* | China/Yunnan 5/Tobacco/1999/Dali | DNA-A: AJ319674 | TYLCCNV/DL |
| *Tomato yellow leaf curl China virus* | China/Yunnan 72/ Datura/2005/Datura | EF011559 | TYLCCNV/Dat |
| *Tomato yellow leaf curl Thailand virus* | Thailand/Chiang Mai/B | DNA-A: AY514630 | TYLCTHV/B |
| *Tomato yellow leaf curl Thailand virus* | China/Yunnan 72/2002/C | DNA-A: AJ495812 | TYLCTHV/C |
| *Tomato yellow leaf curl Thailand virus* | Myanmar/Yangon/1999/D | DNA-A: AF206674 | TYLCTHV/D |
| *Tomato yellow leaf curl Thailand virus* | Thailand/Sakon Nakhon/E | DNA-A: AY514632 | TYLCTHV/E |
| *Tomato yellow leaf curl Guangdong virus* | China/Guangzhou 3/2003 | DNA-A: AY602166 | TYLCGdV |
| *Tomato yellow leaf curl Yunnan virus* | China-YN2013-2011 | DNA-A: KC686705 | TYLCYnV |
| *Tomato yellow leaf curl Mali virus* | Burkina Faso/Tom141/2013 | LM651400 | TYLCMLV |
| tomato yellow leaf curl Mali virus | Ethiopia/Melkassa/2005/Ethiopia | DNA-A: DQ358913 | TYLCMLV/ET |
| *Tomato yellow leaf curl Mali virus* | Mali/2003/Mali | AY502934 | TYLCMLV/ML |
| *Tomato yellow leaf curl Sardinia virus* | Italy/Sardinia/1988 | X61153 | TYLCSaV |
| *Tomato yellow leaf curl Indonesia virus* | Indonesia/Lembang/2005 | DNA-A: AF189018 | TYLCIDV |

**Supplementary table 6. Selected subset of monopartite and bipartite begomoviruses (from Supplementary figure 5).**

| Species | Isolate | Accession number | Virus Abbrev. |
|---|---|---|---|
| *Pepper yellow leaf curl Indonesia virus* | Indonesia/2005 | DNA-A: AB267834; DNA-B: AB267835 | PepYLCIV |
| African cassava mosaic virus | Cameroon | AF112352 | ACMV/CM |

| | | | |
|---|---|---|---|
| *East African cassava mosaic virus* | Uganda/Severe 2/1997/Uganda | DNA-A: AF126806; DNA-B: AF126807 | EACMV/UG |
| *Chilli leaf curl virus* | Pakistan/Multan/1998/Pakistan | DNA-A: AF336806 | ChiLCV/PK |
| *East African cassava mosaic Zanzibar virus* | Tanzania/Uguja/1998 | DNA-A: AF422174; DNA-B: AF422175 | EACMZV |
| *Tomato chlorotic mottle virus* | Brazil/Seabra 1/1996/Bahia | DNA-A: AF490004; DNA-B: AF491306 | ToCMoV/BA |
| *East African cassava mosaic Malawi virus* | Malawi/K/1996 | DNA-A: AJ006460 | EACMMV |
| Tobacco curly shoot virus | isolate Y35 | AJ420318 | TbCSV |
| *Malvastrum yellow vein virus* | China/Yunnan 47/2001 | AJ457824 | MaYVV |
| *Tomato yellow leaf curl virus* | Spain/Almeria/Pepper/1999 | AJ489258 | TYLCV |
| *Tobacco leaf curl Yunnan virus* | China/Yunnan 136/2002/China | AJ512761 | TbLCYnV/CN |
| *Euphorbia leaf curl virus* | China/Guangxi 35/2002 | DNA-A: AJ558121 | EuLCuV |
| *Sri Lankan cassava mosaic virus* | India/Adivaram/2003/India | DNA-A: AJ579307 | SLCMV/IN |
| *Tomato leaf curl Mayotte virus* | Mayotte/Kahani/2003 | AJ865340 | ToLCYTV |
| *Sida yellow mosaic virus* | Brazil/Vicosa 2/1999 | DNA-A: AY090558 | SiYMV |
| *Tomato yellow margin leaf curl virus* | Venezuela/Merida/57 | DNA-A: AY508993; DNA-B: AY508994 | ToYMLCV |
| *Potato yellow mosaic virus* | Venezuela/1991/Potato | DNA-A: D00940; DNA-B: D00941 | PYMV/Po |
| *Mesta yellow vein mosaic virus* | Barackpore | EF373060 | MeYVMV/Ben |
| *Tomato leaf curl Mindanao virus* | Philippines/Mindanao P162/2007 | EU487046 | ToLCMiV |
| *Tomato leaf distortion virus* | Brazil/Paty do Alferes 4/2005 | DNA-A: EU710749 | ToLDV |
| *Tomato yellow leaf distortion virus* | Cuba/5E17/2007 | DNA-A: FJ174698; DNA-B: FJ999999 | ToYLDV |
| *Sunn hemp leaf distortion virus* | India/Barrackpore 1/2008 | DNA-A: FJ455449 | SHLDV |
| *Hollyhock leaf curl virus* | Pakistan/Faisalabad/20/4/06 | FR772082 | HoLCV |
| *Vernonia yellow vein Fujian virus* | CN-Fj-09 | JF265670 | VeYVFV |
| *Tomato yellow mottle virus* | Costa Rica/2003 | DNA-A: KC176780; DNA-B: KC176781 | ToYMoV |
| *Abutilon mosaic virus* | Germany | DNA-A: X15983; DNA-B: X15984 | AbMV |
